# Multiscale Model for Ion Transport in Cellular Media and Applications in Smooth Muscle Cells

**DOI:** 10.1101/2023.04.27.538651

**Authors:** Chun Xiao, Yishui Sun, Huaxiong Huang, Zilong Song, Xingye Yue, Tim David, Shixin Xu

## Abstract

Ion transport in biological tissues is crucial in the study of many biological and pathological problems. Some multi-cellular structures, like the smooth muscles on vessel walls, can be treated as periodic bi-domain structures consisting of the intracellular space (ICS) and extracellular space (ECS) with semipermeable membranes in between. In this work, we first use a multi-scale asymptotic method to derive a macroscopic homogenized bidomain model from the microscopic electro-neutral (EN) model with different diffusion coefficients and nonlinear interface conditions. Then, the obtained homogenized model is applied to study ion transportation and micro-circulation in multi-celluar tissues under the impact of agonists, an internal calcium source, and extracellular potassium. Our model serves as a useful bridge between existing ordinary differential equation models and partial differential models that take into consideration spatial variation. On the one hand, numerical results show that ECS variables are almost invariant in the first two scenarios and confirm the validity of existing single-domain models, which treat variables in the ECS as constants. On the other hand, only the bidomain model is applicable to consider the effect of local extracellular potassium. Finally, the membrane potential of syncytia formed by connected cells is found to play an important role in the propagation of oscillation from the stimulus region to the non-stimulus region.

**Author summary:** Smooth muscle cells (SMCs) play a vital role in neurovascular coupling, which is the mechanism by which changes in neural activity are linked to alterations in blood flow. Dysfunctional SMCs can have significant implications for human health. The activation of SMCs is primarily regulated by the intracellular concentration of calcium ions (Ca2+). A multi-scale model for ion transport in multicellular tissue with varying connectivity has been proposed to investigate SMC activation under different stimuli. The simulation results confirm the critical role of gap junctions in wave propagation and vasoconstriction in the vessel wall. The blockage of gap junctions prevents the spread of the wave. Furthermore, the propagation of membrane potential is the primary cause of wave propagation.

## Introduction

### NVC and SMCs

Neurovascular coupling is the process by which changes in neural activity are coupled to changes in blood flow, which is essential for maintaining proper brain function. When neurons become active and require more energy, they release chemical signals that dilate nearby blood vessels, increasing blood flow to the region and providing the necessary oxygen and glucose for neurons to function.

Smooth muscle cells (SMCs) play a crucial role in neurovascular coupling by regulating the diameter of arterioles, small blood vessels that control blood flow to capillary beds within the brain. Dysfunction of SMCs has been linked to various diseases such as stroke and diabetes [1–3]. Smooth muscle cell (SMC) activation and subsequent contraction of blood vessels is primarily driven by the phosphorylation of actin-myosin motors, which is mainly catalyzed by *Ca*^2+^. There are three different *Ca*^2+^ activation mechanisms reported [1],

- the membrane depolarization due to external stimuli, like electric stimulus, high *K*^+^ concentration;
- internal *Ca*^2+^ release from internal organelles, like the endoplasmic reticulum (ER) or mitochondrial;
- currents that spread through the gap junctions (or connexins) to provide the depolarizing signal;

where the first two mechanisms are related to local activation and the last one induced the propagation of *Ca*^2+^ waves and vasoconstriction in the vessels wall. The propagation plays an important role in coordinating smooth muscle cell activity and facilitating vasoconstriction, which is critical for maintaining proper blood pressure and blood flow.

### Connectivity of Cellular Media

The propagation of waves heavily depends on the connectivity of cellular media. In general, tissues are mainly formed by two compartments, extracellular space (ECS) and intracellular space (ICS). The narrow ECS has filled with an ionic solution in “free diffusion” with the plasma outside cells. The ICS consists of plenty of cells with membranes separating two compartments. On the cell membrane, there are plenty of channels and pumps that allow the passive and active transport respectively of various biological compounds across membrane barriers. On most tissue cells’ membranes, there are gap junctions [4] form conduits between adjacent cells such that the ICS becomes a syncytium for electric wave propagation and ions diffusion. It plays an important role in micro-circulation in tissues [5, 6]. In this case, ECS-ICS forms a “connected-connected” bidomain. However, it is also reported that skeletal muscle does not have any cell-cell junctions [7]. There are no direct communications between cells. In this case, ECS-ICS forms a “connected-disconnected” bidomain. The first goal of this paper is to propose multiscale models of ion transfer in cellular media with these two different spatial connectivities under consideration.

### Models for Ion Transport

The Poisson-Nernst-Planck (PNP) system is one of the most popular mathematical models that describe ion transport under the influence of both ionic concentration gradient and electric field. PNP system has extensive and successful applications in biological systems, particularly in ion channels on the cell membrane [8–13]. However, usually the scale of the bio-tissue we study is much bigger than the scale of a single cell, and this scale separation can cause some difficulty in numerical simulation. Studying ion transport in a multicellular medium requires significant computational resources to carry out a direct simulation [14, 15]. Several approaches are proposed to parallel computing [14], volume average [5, 16–18] and Smeared Multiscale Finite Element Models [19].

There are two main challenges for simulating ion transport in multicellular tissue with PNP equations. First, due to the capacitance of membranes, there are thin boundary layers (BLs) near the interfaces formed by excessive charge accumulation. BLs require extra computation cost during numerical simulations in order to resolve rapid change behaviors of solution inside the layers and attain certain accuracy [20, 21]. Second, due to the existence of pumps on membranes, ion concentrations across the membrane are discontinuous. However, the flux across the membrane is continuous and determined by conductance and the difference between membrane potential and Nernst Potential.

Z. Song et.al. [22] derived effective boundary conditions on the membrane for a one-dimensional EN PNP system and then extended later to higher dimensions [23]. Similar effective boundary conditions were derived in [24] where the Poisson equation is replaced by the EN condition. In [25], by taking linearization of the effective boundary conditions in [23], a microscopic EN bi-domain ion transport model with membrane capacity interface term is proposed, and the corresponding homogenized system is derived rigorously by unfolding operator [26, 27] in homogenization theory [28–34], with the assumption that the diffusion coefficients are identical for all ion species. In this work, we extended to the case when the diffusion coefficients are different with a more general form of interface flux. Multiscale expansion [35] is used to derive the macroscopic homogenization system for both the ICS part and the ECS part.

Then the proposed multiscale model is applied to investigate ions transport in vascular smooth muscle cells (VSMCs) with different internal and external stimuli. With our bidomain model, we obtain three main results. Firstly, the critical points of ICS calcium source, PLC and ECS potassium concentration are identified. Secondly, the bidomain model confirms that the ECS variation is negligible when ICS calcium concentration oscillates due to variations in ICS calcium and PLC. Thirdly, the spatial wave propagations from the stimulus region to the non-stimulus region are mainly induced by the membrane potential of syncytia formed by connected cells rather than the ions.

## Mathematical Model

### Microscale model

In this section, the microscale model for ion transport is proposed. As shown in Figure 1 (a), the multi-celluar domain Ω_*L*_ = (0, *L*)^*d*^ consists of the ICS (Ω_*I*_) and ECS (Ω_*E*_). Γ_*m*_ = ∂Ω_*I*_ is the interface of ICE and ECS formed by the cells’ membranes. For simplicity, we assume Ω_*L*_ to be a union of *Y*_*l*_-periodic sets with *Y*_*l*_ = (0, *l*)^*d*^ and *L/l* ∈ ℕ. The narrow ECS Ω_*E*_ is very tortuous but fully connected. While for ICS Ω_*I*_, in this work, both connected or unconnected cases are considered. Figure 1(a) only shows the case when Ω_*I*_ is not connected.

**Fig 1.**
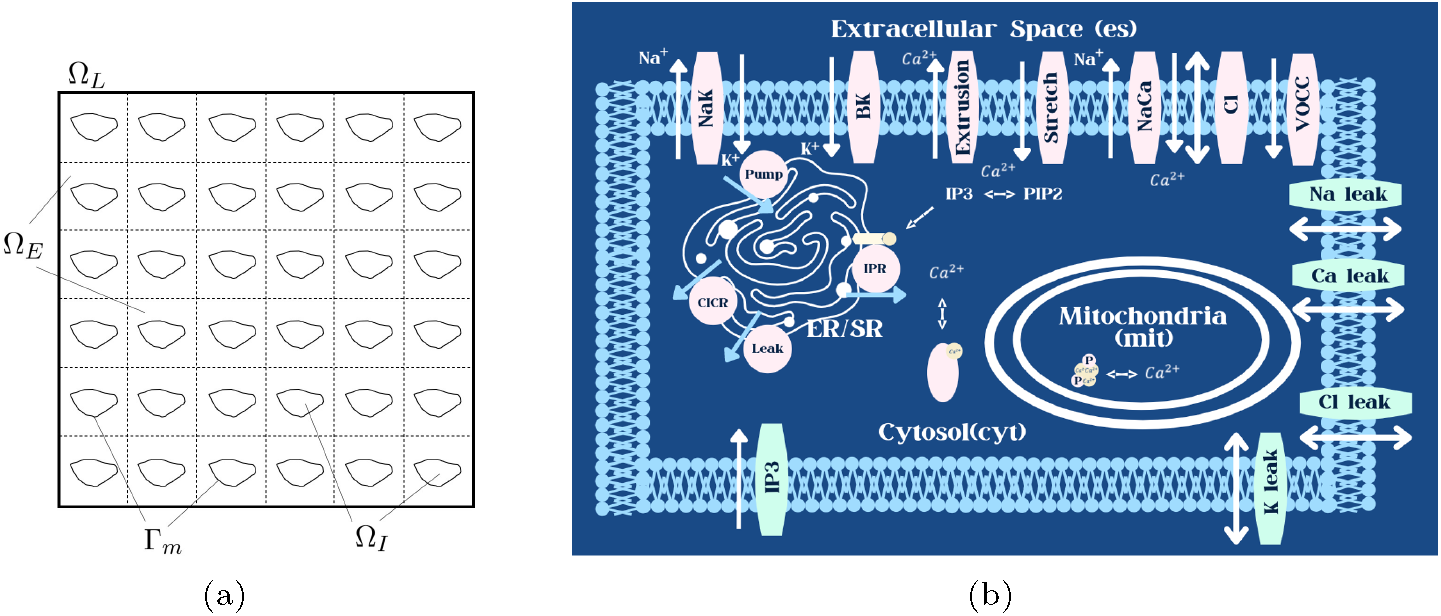
(a): Schematic of the domain Ω_*L*_; (b): Schematic diagram of the SMC model containing *IP*_3_, K^+^, Cl^−^, Na^+^ and Ca^2+^ ions. The SMC contains the SR acting as an internal store of Ca^2+^. There are eleven channels on the cell membrane and four channels on the SR membrane.

Suppose there are *N* ionic species under consideration. Their concentrations and valences are *C*_*i,s*_ *>* 0 and *z*_*i*_, *i* = 1, 2, …, *N*, respectively; subscript *s* stands for variables in Ω_*s*_, *s* = *I, E*. Since the microscale is the cell scale (micrometer) which is much larger than the Debye length, the following EN assumption is imposed [22, 23]:

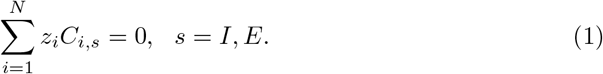

The microscopic EN bi-domain model for ion concentration is given as follows, for *i* = 1, …, *N, s* = *I, E*,

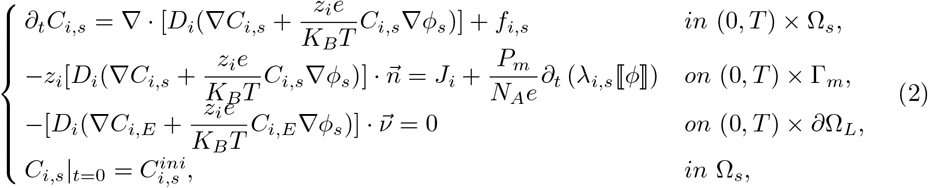

where *ϕ*_*s*_ is the electric potential in Ω_*s*_, *D*_*i*_ is constant diffusion coefficient for the i*th* ion, *f*_*i,s*_ is the source term, 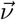 is the outer boundary normal direction, 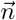 is the normal direction of cell membrane, pointing from ICS to ECS. ⟦*ϕ*⟧ = *ϕ*_*I*_ − *ϕ*_*E*_ is the membrane potential. On the membrane, the ions’ fluxex consist of two parts, the one induced by channels on the cell membrane and the other induced by the capacitance effect of membrane. The former is usually a function of ⟦*ϕ*⟧ and *C*_*i,s*_, i.e.,

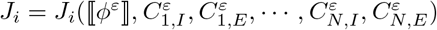. The latter has a parameter 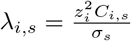 with 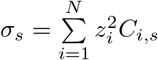 the ratio of capacitance effect on the i*th* ion [22, 23]. *P*_*m*_ is the membrane capacitance, *k*_*B*_ is the Boltzmann constant, *T* is temperature, *N*_*A*_ is the Avogadro constant. The values are shown in Table 1.

**Table 1.**
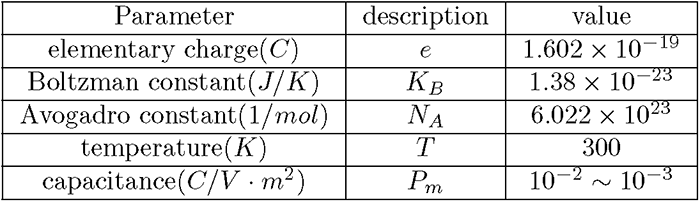
Parameters

| Parameter | description | value |
| --- | --- | --- |
| elementary charge($C$) | $e$ | $1.602 \times 10^{-19}$ |
| Boltzman constant($J/K$) | $K_B$ | $1.38 \times 10^{-23}$ |
| Avogadro constant($1/mol$) | $N_A$ | $6.022 \times 10^{23}$ |
| temperature($K$) | $T$ | 300 |
| capacitance($C/V \cdot m^2$) | $P_m$ | $10^{-2} \sim 10^{-3}$ |

Multiplying concentration equations with *z*_*i*_ and summing up with EN condition (1) and (2) yield the equation for electric potential:

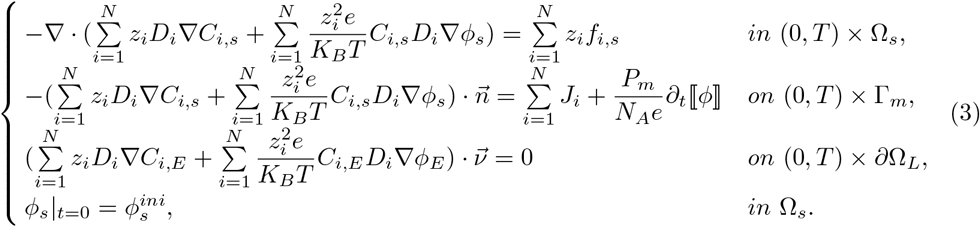

Before carrying out the homogenization, we first implement the following nondimensionalization:

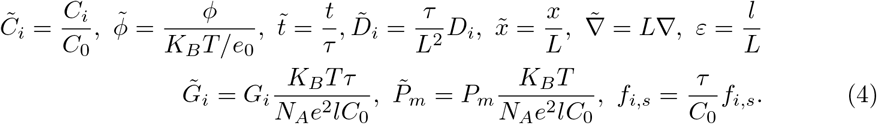

The scale of ion concentration we consider here ranges from 10^−4^ *mM* to 10^2^ *mM*, so the reference concentration is set to be *C*_0_ = 1 *mM*. The reference time is *τ* = 1 *s* since the time scale in our numerical simulation is about 10^2^ *s. L* = 10^−3^ *m* is the reference macro scale, *l* = 13 *μm* is the reference micro scale.

Then the dimensionless region is Ω = (0, 1)^*d*^ which consists of two components: 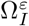 and 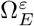. Let *Y* = (0, 1)^*d*^ and *Y*_*I*_, *Y*_*E*_ are two disjoint subsets of *Y*, such that

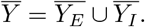

And Γ = ∂*Y*_*I*_ ⋂ ∂*Y*_*E*_ is smooth. For any *k* ∈ ℤ^*d*^, let

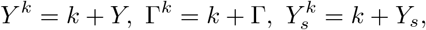

where *k* = (*k*_1_, …, *k*_*d*_), *s* = *I, E*. For any *ε >* 0 and 1*/ε* ∈ ℕ^+^, let 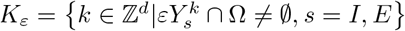. Denote the two disjoint subsets of Ω and the interface between them as:

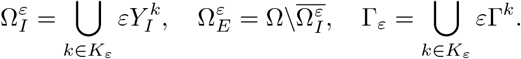

Denote the two disjoint subsets of Ω and the interface between them as:

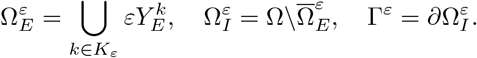

Note that ∂Ω and Γ_*ε*_ are disjoint, 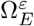 is connected. Based on the connectivity of 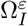, we will consider both “connected-disconnected” and “connected-connected” cases.

The dimensionless equations for (2) and (3) are

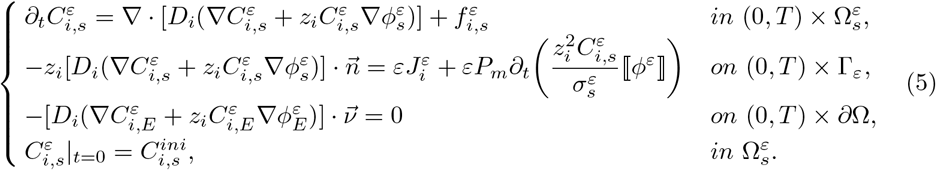

and

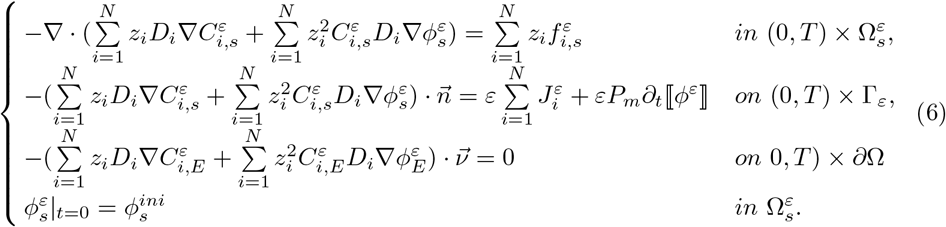

where 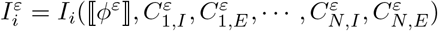 and ∼ is omitted for simplicity.

### Homogenized macro-scale model

In this section the homogenized equations for (5) and (6) are derived using asymptotic expansion. The main results are presented in the following theory. The detailed derivation are given in Appendix 0.0.1.

#### Theorem 0.1.

*Suppose* 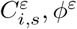, *ϕ*^*ε*^ *are the solutions of* (5),(6), *and* 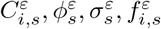 *have formal two-scale asymptotic expansions of the form*

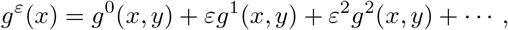

*where g are Y* −*periodic with respect to y* = *x/ε and* 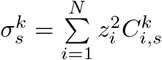 *for k* = 0, 1, 2, … .*Suppose I*_*i*_ *is smooth, then* 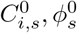 *are independent of y and the following macro models hold*.

1. *In* “*connected-disconnected*” *case*, 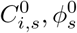 *satisfy the following upscaled system:*

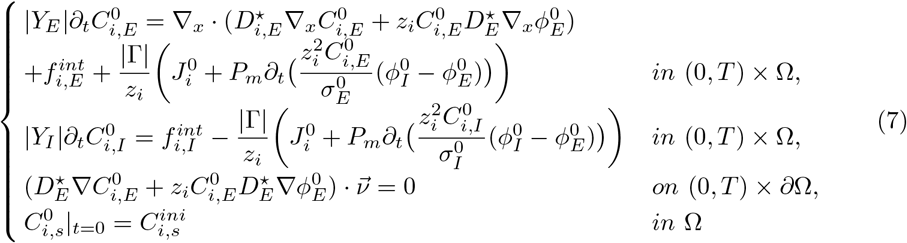

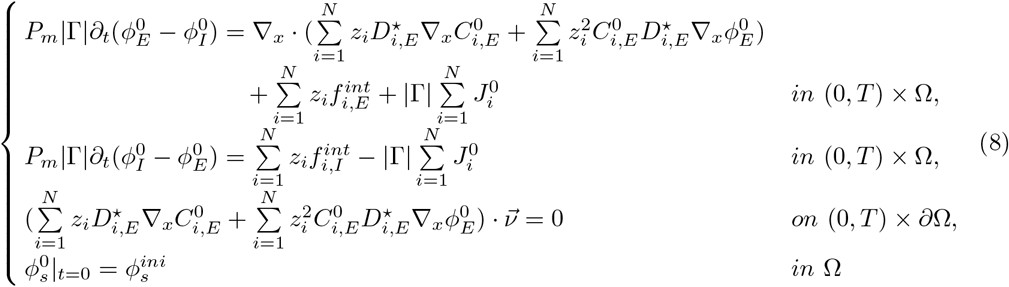
2. *In* “*connected-connected*” *case*, 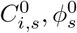 *satisfy the following upscaled system:*

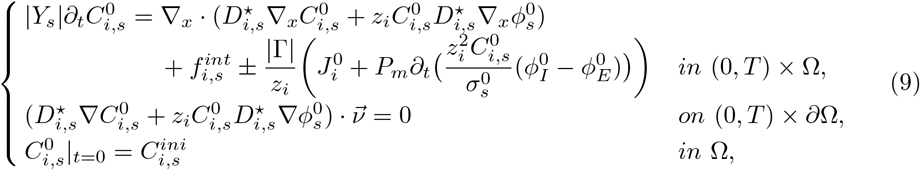

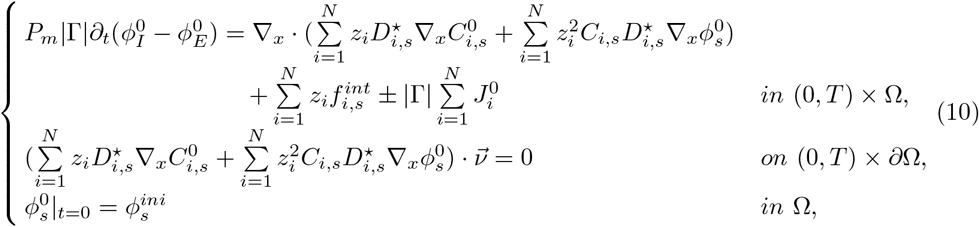

*for j* = 1, …, *d and*”*±*”*is* “+” *for s* = *E and* “−” *for s* = *I. And* 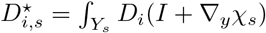 *dy, satisfy*

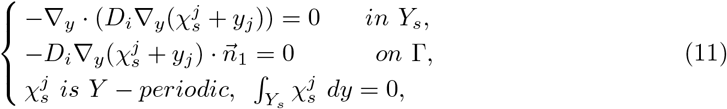

*for s* = *I, E*. 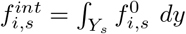.

#### Remark 0.1.

*The theorem shows that after upscaling, the transmembrane fluxes in the micro-scale are changed to be source/sink terms in the bulk equations to describe the micro-circulation between ICS and ECS. For the* “*connected-connected*” *case, the transport of ions in two compartments are described by diffusion-convection equations with a source/sink term due to the transmembrane communication between two compartments. While for the* “*connected-disconnected*” *case, the dynamics of each ion concentration is an ordinary differential equation inside the ICE. Also, when the micro-scale diffusion coefficient in connexin region D*_*B*_ *decays to zero, the* “ *connected-connected*” *is degenerated to the* “*connected-disconnected*” *case, as the* 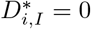.

### Model for smooth muscle cells

Finally, homogenized equations (9) and (10) is used to study the ion transport in VSMCs. Here four types of ions, calcium, chloride, potassium, sodium are taken into consideration denoted by [*K*^+^]_*s*_, [*Cl*^−^]_*s*_, [*Na*^+^]_*s*_, [*Ca*^2+^]_*s*_, with *s* = *I, E* for ICS and ECS, respectively. As shown in Fig. 1b, in side the cell, the communication between endoplasmic reticulum (ER)/(SR) with calcium denoted by [*Ca*^2+^]_*SR*_ and cytosol are taken into consideration to study the effect of IP3. We also included a source term induced by the nucleartion of calcium in mitochondria [36]. On the VSMC’s membrane, there are pumps and channels for active and passive transmembrane flux. Na-K is a *Na*^+^*/K*^+^ ATPase pump which pumps three *Na*^+^ out of the cell and two *K*^+^ into the cell [37]. BK is the voltage-gated big potassium channel that conduct large amounts of *K*^+^ across the cell membrane [38]. Extrusion channel represent the *Ca*^2+^ extrusion from the SMC by *Ca*^2+^ ATPase pump [39]. Stretch channel can induce release of *Ca*^2+^ from ICS via activated second messenger systems [40]. Na-Ca is an *Na*^+^*/Ca*^2+^ exchanger which transfer three *Na*^+^ into the cell and one *Ca*^2+^ out of the cell [41]. [*Ca*^2+^]_*I*_ is increased through activation of the calcium-activated Cl channel (chloride channel) after agonist stimulation [41]. The *IP*_3_ represents the effect that G-protein-coupled receptors induce the activation of Phospholipase C (PLC), which cause *IP*_3_ diffuse into the cytosol. The four leak channels on the cell membrane represent other fluxes which are not considered in our model. For the ionic fluxes on the membrane of SR, firstly, *IP*_3_ can induce SR *Ca*^2+^ flux into the cell cytosol, and it would enhance the flux of *Ca*^2+^ out of the SR reticulum [42], this process is represented by CICR channel. IPR channel denote *IP*_3_-induced SR *Ca*^2+^ flux into ICS. *Ca*^2+^ leak from SR is denoted by leak channel. Pump channel utilize energy from ATP to pump *Ca*^2+^ back to the SR against a concentration gradient [41]. The detailed expression of these fluxes could be found in Appendix 0.0.1. The parameters in (12), (14), (13) are in Table 2.

**Table 2.** Model parameters

| Parameter | Description | dimensional value | dimensionless value | note |
| --- | --- | --- | --- | --- |
| $\vert \Gamma\vert $ | Ratio of the surface area to the volume of the cell | 5.9653 | 5.9653 | this work |
| $z_1$ | Ionic valence for $\text{Ca}^{2+}$ | 2 | 2 | |
| $z_2$ | Ionic valence for $\text{K}^{2+}$ | 1 | 1 | |
| $z_3$ | Ionic valence for $\text{Cl}^{2+}$ | -1 | -1 | |
| $z_4$ | Ionic valence for $\text{Na}^{2+}$ | 1 | 1 | |
| $\vert Y_I\vert $ | volume fraction of ICS | 0.75 | 0.75 | [43] |
| $\vert Y_E\vert $ | volume fraction of ECS | 0.25 | 0.25 | [43] |
| $\lambda$ | Rate constant for opening the BK channel | $45\text{s}^{-1}$ | 45 | this work |
| $C_i$ | CICR rate constant | $0.009\text{ mM s}^{-1}$ | 0.009 | this work |
| $s_c$ | Half-point of the CICR $\text{Ca}^{2+}$ efflux sigmoidal | $53.33 \times 10^{-3}\text{ mM}$ | $53.33 \times 10^{-3}$ | this work |
| $c_c$ | Half-point of the CICR activation sigmoidal | $9 \times 10^{-4}\text{ mM}$ | $9 \times 10^{-4}$ | this work |
| $B_i$ | SR uptake pump rate constant | $2.0 \times 10^{-3}\text{ mM s}^{-1}$ | $2.0 \times 10^{-3}$ | this work |
| $c_b$ | Half-point of the SR uptake pump activation sigmoidal | $1 \times 10^{-3}\text{ mM}$ | $1 \times 10^{-3}$ | this work |
| $L_i$ | Leak from SR rate constant | $0.0012\text{ s}^{-1}$ | 0.0012 | this work |
| $V_{M3}$ | Maximal value for $\text{Ca}^{2+}$ release from SR | $0.1\text{ mM s}^{-1}$ | 0.1 | this work |
| $K_a$ | $[\text{Ca}^{2+}]_I$ dependent threshold constant for $J_{SR,IPR}$ | $0.9 \times 10^{-3}\text{ mM}$ | $0.9 \times 10^{-3}$ | this work |
| $K_r$ | $[\text{Ca}^{2+}]_{SR}$ dependent threshold constant for $J_{SR,IPR}$ | $53.33 \times 10^{-3}\text{ mM}$ | $53.33 \times 10^{-3}$ | this work |
| $K_p$ | $\text{IP}_3$ dependent threshold constant for $J_{SR,IPR}$ | $0.65 \times 10^{-3}\text{ mM}$ | $0.65 \times 10^{-3}$ | this work |
| $F_{NaK}$ | Rate of $\text{K}^+$ influx by the $\text{Na}^+/\text{K}^+$ pump | $2.171 \times 10^{-4}\text{ C m}^{-2}\text{ s}^{-1}$ | $4.325 \times 10^{-5}$ | this work |
| $G_{VOCC}$ | Conductance of the VOCC | $0.0338\text{ CV}^{-1}\text{ m}^{-2}\text{ s}^{-1}$ | $2.43 \times 10^{-4}$ | this work |
| $v_{Ca2}$ | Half-point of the VOCC activation sigmoidal | $-0.024\text{ V}$ | $-0.928$ | this work |
| $\Delta p$ | Pressure difference over vessel | $30\text{ mmHg}$ | 30 | this work |
| $P_m$ | capacitance | $0.0019\text{ C}/(\text{V} \cdot \text{m}^2)$ | $1 \times 10^{-5}$ | this work |
| $\alpha$ | volume fraction of SR | 0.05 | 0.05 | [41] |
| $R_{Ca}$ | Maximum slope of the VOCC activation sigmoidal | $8.5 \times 10^{-3}\text{ V}$ | 0.329 | this work |
| $D_i$ | Rate constant for $\text{Ca}^{2+}$ extrusion by the adenosine triphosphatase (ATP)-ase pump | $0.801\text{ CV}^{-1}\text{ m}^{-2}\text{ mM}^{-1}\text{ s}^{-1}$ | 0.166 | this work |
| $v_d$ | Intercept of voltage dependence of extrusion ATP-ase | $-0.1\text{ V}$ | $-3.868$ | this work |
| $R_d$ | Slope of voltage dependence of extrusion ATP-ase | $0.25\text{ V}$ | 9.67 | this work |
| $G_{stretch}$ | Conductance of the $\text{Ca}^{2+}$ stretch channel | $0.0062\text{ CV}^{-1}\text{ m}^{-2}\text{ s}^{-1}$ | $3.295 \times 10^{-5}$ | this work |
| $\alpha_{stretch}$ | Slope of stress dependence of the stretch activation sigmoidal | $0.0074\text{ mmHg}^{-1}$ | 0.0074 | this work |
| $R/h$ | Ratio of the vessel radius to the vessel wall thickness | 10 | 10 | this work |
| $\sigma_0$ | Half-point of the stretch activated channel (SAC) activation sigmoidal | $500\text{ mmHg}$ | 500 | this work |
| $G_{NaCa}$ | Conductance of the $\text{Na}^+/\text{Ca}^{2+}$ exchanger | $0.0049\text{ CV}^{-1}\text{ m}^{-2}\text{ s}^{-1}$ | $2.62 \times 10^{-5}$ | this work |
| $G_{ClCa}$ | Conductance of the $\text{Cl}^-$ channel | $0.0196\text{ CV}^{-1}\text{ m}^{-2}\text{ s}^{-1}$ | $1.0486 \times 10^{-4}$ | this work |
| $k_{CACC}$ | dissociation constant of CACC | $0.587 \times 10^{-3}\text{ mM}$ | $0.587 \times 10^{-3}$ | this work |
| $J_{PLC}$ | $\text{IP}_3$ production rate | $0\text{ to }10^{-3}\text{ mM s}^{-1}$ | $0\text{ to }10^{-3}$ | this work |
| $k_d$ | Rate constant of $\text{IP}_3$ degradation | $0.1\text{ s}^{-1}$ | 0.1 | this work |
| $\alpha_{act}$ | Translation factor for voltage dependence of $K_{act_i}$ sigmoidal | $1.32 \times 10^{-7}\text{ mM}^2$ | $1.32 \times 10^{-7}$ | this work |
| $v_{Ca3}$ | Half-point for the $K_{act_i}$ activation sigmoidal | $-0.027\text{ V}$ | $-1.044$ | this work |
| $R_k$ | Maximum slope of the $K_{act_i}$ activation sigmoidal | $0.012\text{ V}$ | 0.464 | this work |
| $G_{BK}$ | Conductance of the BK channel | $0.011\text{ CV}^{-1}\text{ m}^{-2}\text{ s}^{-1}$ | $5.878 \times 10^{-5}$ | this work |
| $v_{SAC}$ | Nernst potential of the stretch channel | $0.1034\text{ V}$ | 4 | this work |
| $G_{Ca,leak}$ | Conductance of calcium leak channel | $1.51 \times 10^{-6}\text{ CV}^{-1}\text{ m}^{-2}\text{ s}^{-1}$ | $8.08 \times 10^{-9}$ | this work |
| $G_{K,leak}$ | Conductance of potassium leak channel | $0.017 \times 10^{-2}\text{ CV}^{-1}\text{ m}^{-2}\text{ s}^{-1}$ | $9.28 \times 10^{-5}$ | this work |
| $G_{Cl,leak}$ | Conductance of chloride leak channel | $-3.2 \times 10^{-6}\text{ CV}^{-1}\text{ m}^{-2}\text{ s}^{-1}$ | $-1.69 \times 10^{-5}$ | this work |
| $G_{Na,leak}$ | Conductance of sodium leak channel | $3.9 \times 10^{-3}\text{ CV}^{-1}\text{ m}^{-2}\text{ s}^{-1}$ | $2.10 \times 10^{-5}$ | this work |

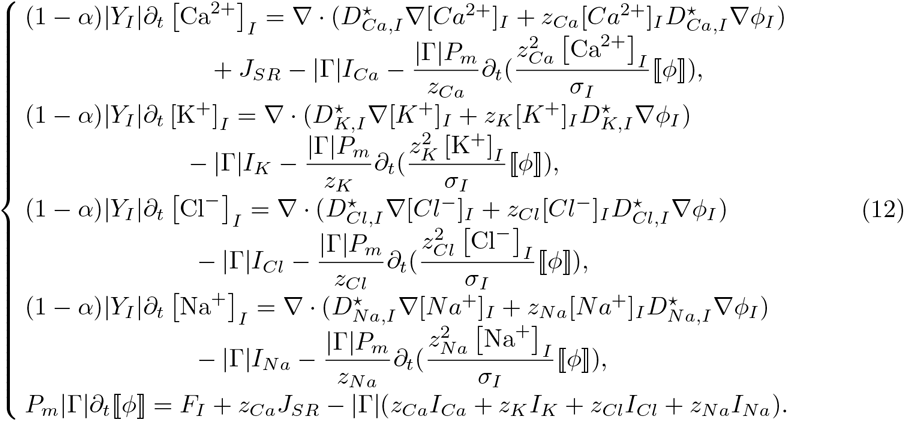

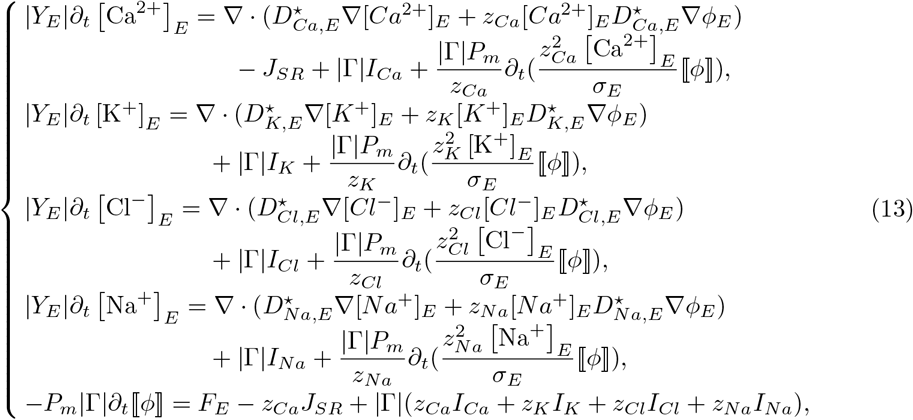

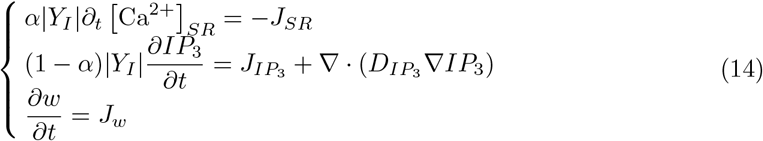

where *w* is the open probability of BK channel, |*Y*_*I*_| and |*Y*_*E*_| are the volume fraction of *Y*_*I*_ and *Y*_*E*_ in *Y, α* is the volume fraction of ER/SR in VSMC, and

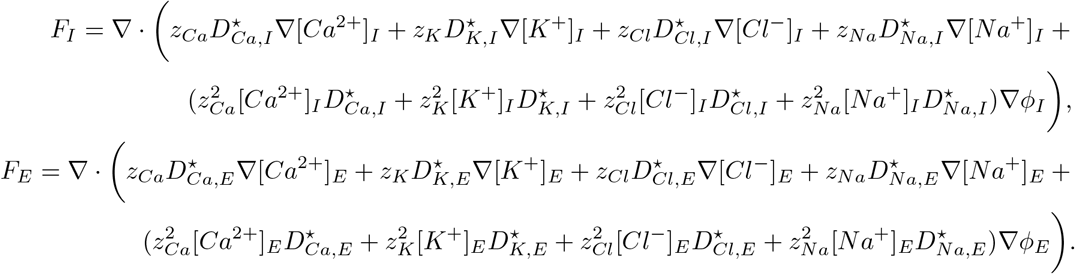

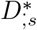 are the effective diffusion coefficients and the derivation is shown in Appendix 0.0.1. The initial values for (12), (13), (14) are in Table 3. Here the symbol (·)^0^ is omitted for simplicity.

**Table 3.**
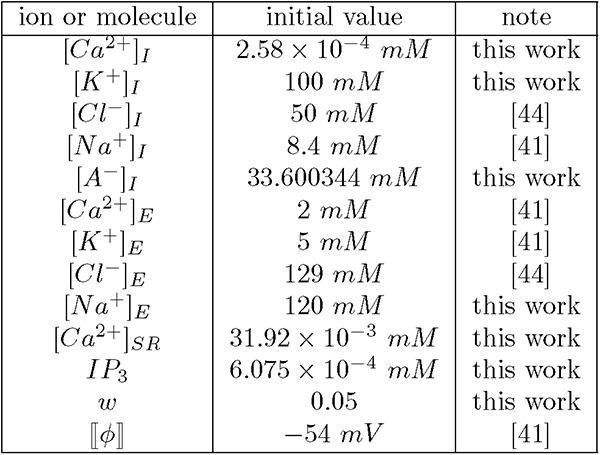
Initial value

| ion or molecule | initial value | note |
| --- | --- | --- |
| $[Ca^{2+}]_I$ | $2.58 \times 10^{-4} \text{ mM}$ | this work |
| $[K^+]_I$ | $100 \text{ mM}$ | this work |
| $[Cl^-]_I$ | $50 \text{ mM}$ | [44] |
| $[Na^+]_I$ | $8.4 \text{ mM}$ | [41] |
| $[A^-]_I$ | $33.600344 \text{ mM}$ | this work |
| $[Ca^{2+}]_E$ | $2 \text{ mM}$ | [41] |
| $[K^+]_E$ | $5 \text{ mM}$ | [41] |
| $[Cl^-]_E$ | $129 \text{ mM}$ | [44] |
| $[Na^+]_E$ | $120 \text{ mM}$ | this work |
| $[Ca^{2+}]_{SR}$ | $31.92 \times 10^{-3} \text{ mM}$ | this work |
| $IP_3$ | $6.075 \times 10^{-4} \text{ mM}$ | this work |
| $w$ | $0.05$ | this work |
| $[\phi]$ | $-54 \text{ mV}$ | [41] |

#### Remark 0.2.

*Note that we add a source term z*_*Ca*_*J*_*SR*_ *in the equation for Cl*^−^ *in* (13) *due to the charge neutrality*.

## Simulation Results and Discussion

### Model response to varying Phospholipase C

The SR mainly serves as an intracellular calcium storage compartment and plays an important roles on calcium-release-dependent cellular processes through different channels on its membrane. One of the most channels for Calcium leaking from SR to cytosol is IPR channel. Its open probability is regulated by cytosolic calcium concentration and IP3 concentration, which is produced by PLC [41].

#### Single cell dynamics

Initially we test the model with spatially homogeneous initial conditions (ICS) and no flux boundary conditions on ∂Ω. If the variation of *J*_*PLC*_ is also homogeneous in the space, the dynamics of the VSMC behaves like a one single cell.

Figure 2 shows the results of intracellular ions and membrane potential ⟦*ϕ*⟧ when the *J*_*PLC*_ variates from the default value 0.243 *μMs*^−1^ (blue line) to *J*_*PLC*_ = 0.4 *μMs*^−1^(red lines). When the *J*_*PLC*_ increases, the *IP*_3_ increases and achieves a new equilibrium. The intracelluar *Ca*^2+^ concentrations in cytosol and SR, membrane potential ⟦*ϕ*⟧ are BK open probability all oscillate within a physiological range. The *K*^+^, *Na*^+^ and *Cl*^−^ concentrations do not oscillate with any significant amplitude. The results in Appendix Figure 15 show that the ions in the ECS do not have observable oscillation when more PLC is supplied. This confirms that the rationality of existing models assumptions on constant ECS variables.

**Fig 2.**
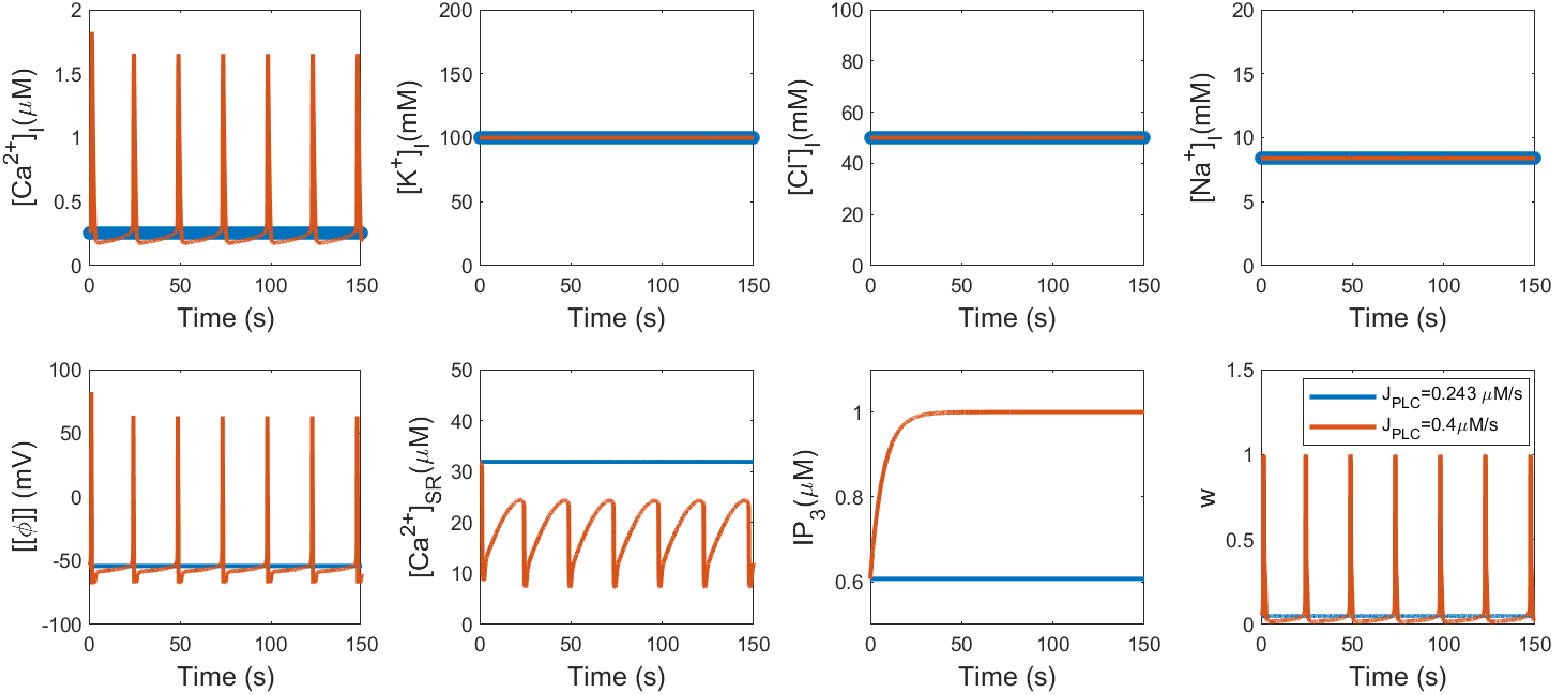
Intracellular variable dynamics for the case of space homogeneous problem: Blue lines: *J*_*P LC*_ = 0.243 *μMs*^−1^ (Resting state); Red lines: *J*_*P LC*_ = 0.4 *μMs*^−1^.

##### Remark 0.3.

*From Figure 2 and Figure 15 in Appendix, it can be seen that when J*_*PLC*_ *is high*, [*Ca*^2+^]_*I*_ *is oscillating and* [*Ca*^2+^]_*E*_ *is not. Actually*, [*Ca*^2+^]_*E*_ *is oscillating at the same scale as* [*Ca*^2+^]_*I*_ *is, but the amplitude is too small to be seen since* [*Ca*^2+^]_*E*_ *is several scale higher than* [*Ca*^2+^]_*I*_. *The same can be said for* [*Ca*^2+^]_*E*_ *and* [*K*^+^]_*E*_ *in the following sections*.

Figure 3(a) showed the bifurcation diagram of [*Ca*^2+^]_*I*_ as *J*_*PLC*_ vary continuously from 0 *μMs*^−1^ to 0.5 *μMs*^−1^. The critical value of *J*_*PLC*_ for [*Ca*^2+^]_*I*_ to oscillate is 0.243 *μMs*^−1^. The reuslt in Figure 3(b) illustrated that the oscillation frequency increases with the *J*_*PLC*_. In Panel (c), it shows when *J*_*PLC*_ is high, the IPR channel flux of *Ca*^2+^ from SR to cytosol induced the elevation of [*Ca*^2+^]_*I*_ and decrease of [*Ca*^2+^]_*SR*_, which is the main cause of oscillation.

**Fig 3.**
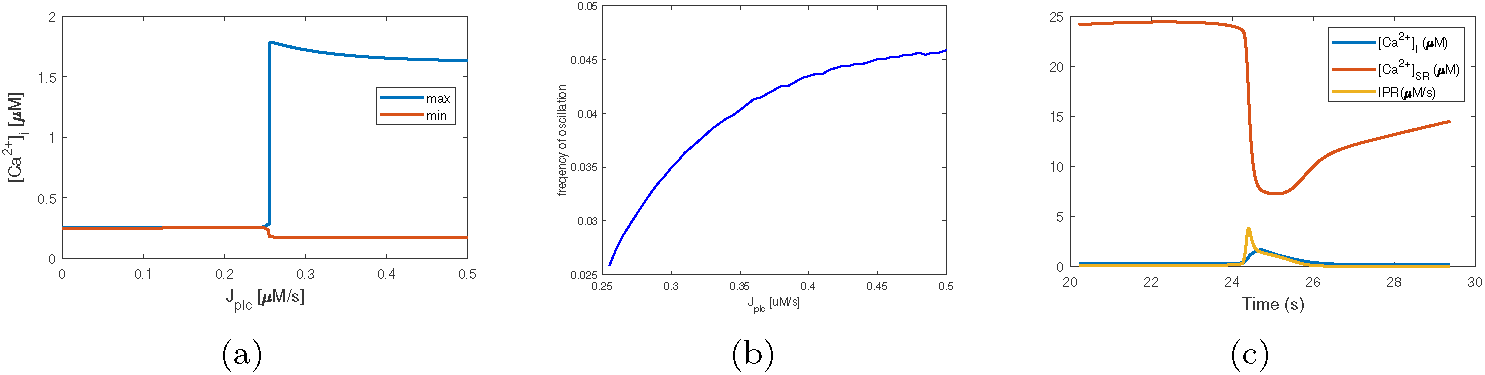
(a) Bifurcation diagram for [*Ca*^2+^]_*I*_ when *J*_*PLC*_ vary from 0 *μMs*^−1^ to 0.5 *μMs*^−1^; (b) Oscillation frequency with different *J*_*PLC*_ ; (c) Flux of *Ca*^2+^ through IPR channel when *J*_*PLC*_ = 0.4 *μMs*^−1^.

#### **Spatially varied** *J*_*PLC*_

In physiology it is unlikely that the ATP concentration (and hence *J*_*PLC*_) would be constant over space; instead there is likely to be some spatial variation over a surface such as an artery due to factors such as wall shear stress (WSS) [15]. In order to model this spatial variation we vary *J*_*PLC*_ linearly over a domain (see Figure 18). In this case, the full spatial model with/without diffusion in Ω is considered.

We consider a long 2-D domain Ω_*r*_ = [0, 100] × [0, 2] with *J*_*PLC*_ vary linearly over *x*_1_ direction where at *x*_1_ = 0, *J*_*PLC*_ = 0 *μ*Ms^−1^ and at *x*_1_ = 100, *J*_*PLC*_ = 0.5 *μ*Ms^−1^. The initial conditions are spatially homogeneous, and non-flux boundary conditions are imposed on each boundary, finite volume method is applied to solve the model.

In Figure 4(a,b,d,e), the dynamics of ICS calcium [*Ca*^2+^]_*I*_ and electric potential ⟦*ϕ*⟧ for a single “row” (*x*_2_ = 1) of Ω_*r*_ is presented for both cases. For the “connected-disconnected” case, the oscillation is only observed in the region *x*_1_ *>* 48 where *J*_*PLC*_ ≥ 0.24. While for the “connected-connected” case, the oscillation is observed in the whole domain. The wave proper gates from high *J*_*PLC*_ region to low *J*_*PLC*_ region.

**Fig 4.**
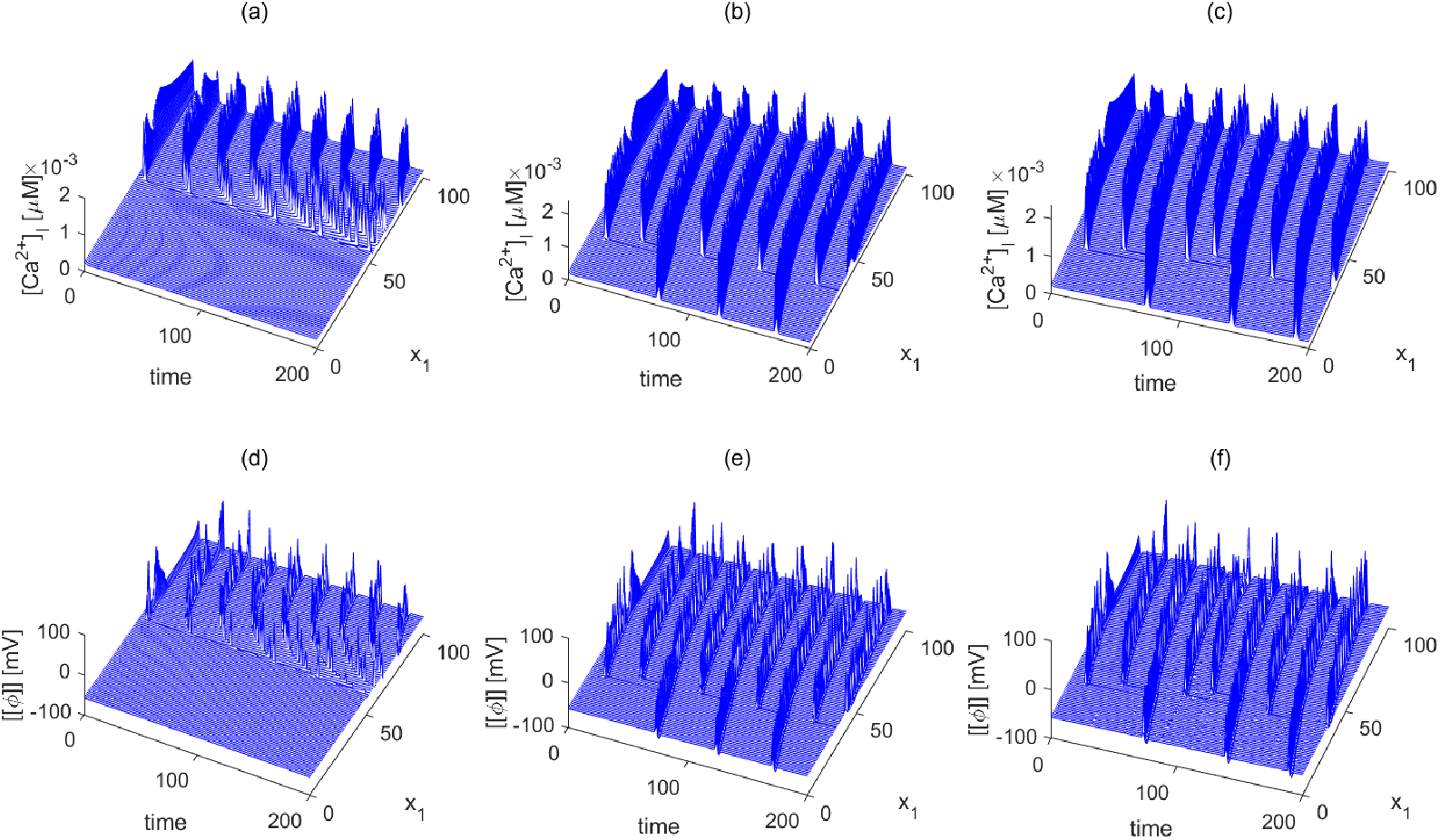
Dynamics of [*Ca*^2+^]_*I*_ and ⟦ *ϕ* ⟧ for a single “row” (*x*_2_ = 1) of Ω_*r*_. (a) and (d): connected-disconnected case; (b) and (e): connected-connected case. (c) and (f): connected-connected case without ion diffusion.

In Figure 5, the effect of gap junctions on wave propagation is further studied [45]. As the the gap junctions are partially blocked, the effective diffusion constant decrease to ten percent of the original one. It shows that the propagation of *Ca*^2+^ wave fade away in the non-stimulus region. This confirms the gap junctions roles in coordinating smooth muscle cell activity and facilitating vasoconstriction.

**Fig 5.**
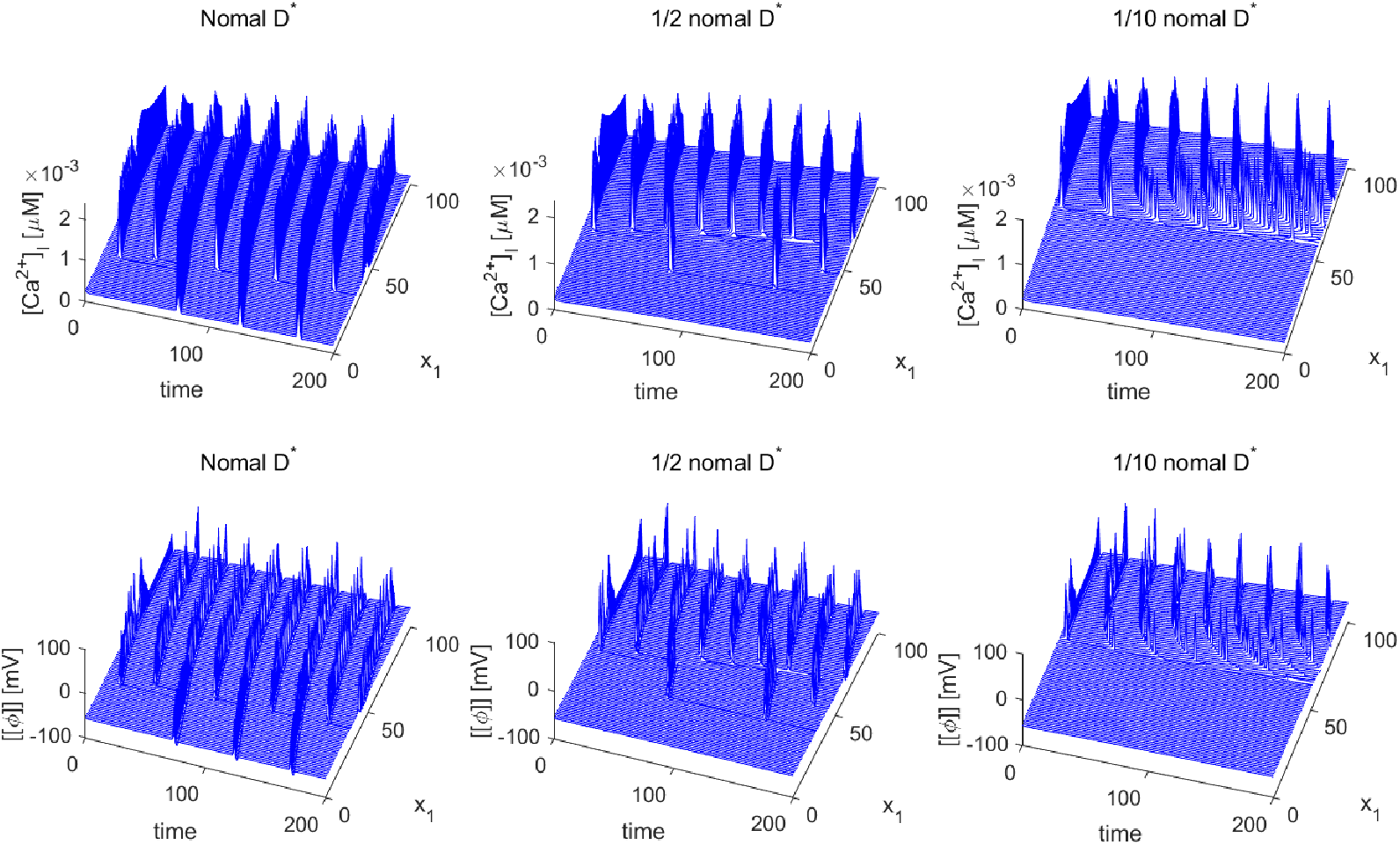
Dynamics of [*Ca*^2+^]_*I*_ and ⟦*ϕ*⟧ for a single “row” (*x*_2_ = 1) of Ω_*r*_. As *D*^⋆^ decreases, the propagation of oscillation also diminishes.

### Model response to varying extracellular potassium concentration

In this section, the effects of potassium on the dynamics of calcium and membrane potential are studied. Both Experiments results [46, 47] and simulation results [18, 36] show that [*K*^+^]_*E*_ induced *Ca*^2+^ oscillation can cause contractions of SMCs in intrapulmonary bronchioles and intrapulmonary arterioles.

#### Single cell dynamics

Similarly, We first consider the homogeneous case, when the ECS potassium is elevated uniformly in space. The behaviors of cytosol calcium and membrane potential with different ECS potassium concentration are shown in Figure 6. When the ECS potassium increases from default value 5*mM* (in red) to 16*mM* (in green). The oscillations of ICS and membrane potential are observed. While, keeping increase the ECS potassium to 25.1*mM* (in blue), the oscillation of ICS potassium stops and the membrane potential stops oscillate and shift away from the resting value, which means the depolarization. Appendix Figure 16 confirms that there are no observable variations in the ECS variables.

**Fig 6.**
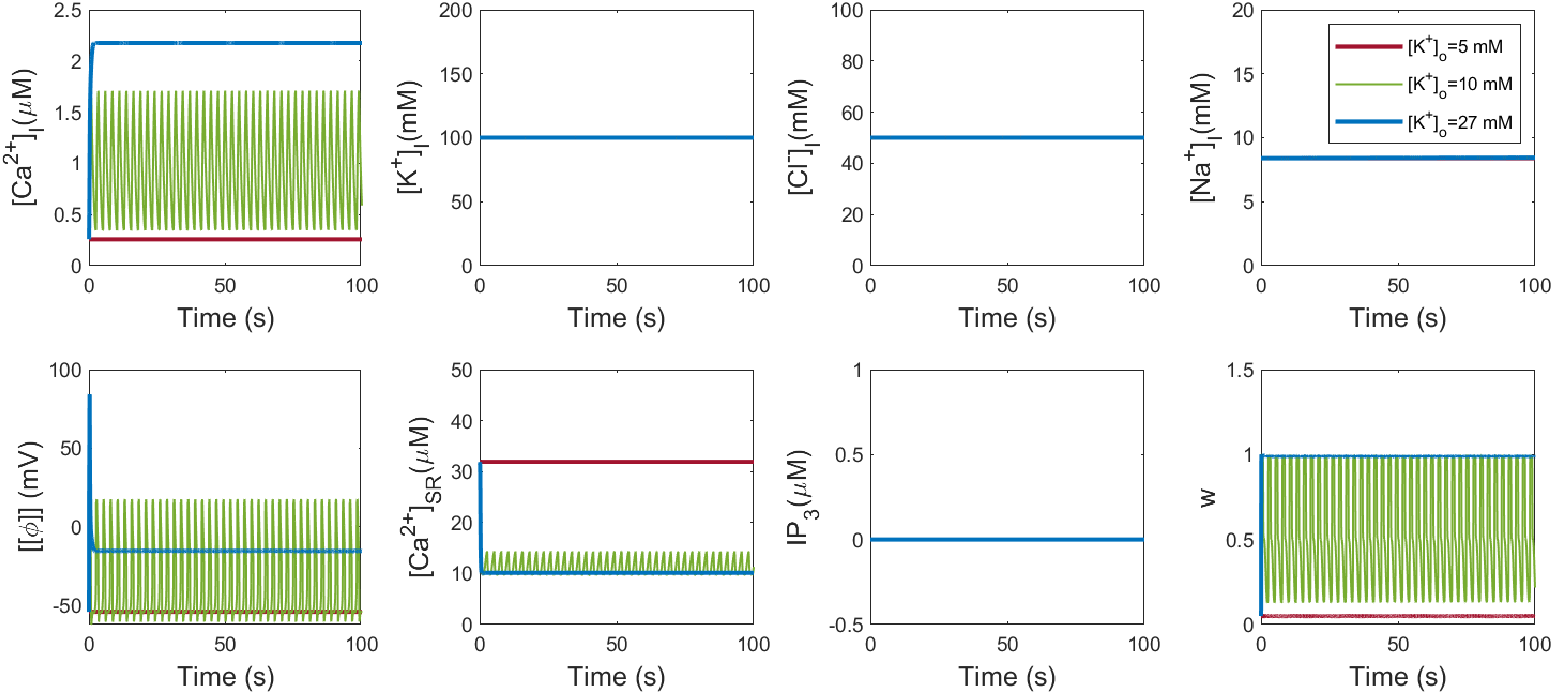
Intracellular variable dynamics as [*K*^+^]_*E*_ increases for the case of space homogenuous problem. Red lines: 5*mM* (Resting state); Green lines: 10*mM* ; Blue lines: 27*mM*.

In order to critical the critical values of [*K*^+^]_*E*_ for [*Ca*^2+^]_*I*_ to begin oscillation and to exhibit depolarization, the Hopf bifurcation diagram is shown in Figure 7. The critical value of [*K*^+^]_*E*_ for inducing depolarization is 25.1 *mM*. The threshold of [*K*^+^]_*E*_ for depolarization is consistent with the experiment in [48], where high resistance capillary micro-electrodes were used to record membrane potentials in an attempt to minimize the degree of cell injury in the smooth muscle of the taenia coli of the guinea pig, and their results showed that the concentration of [*K*^+^]_*E*_ for depolarization is between 23 *mM* and 46 *mM*. From Figure 6 and Figure 7 we can see that as [*K*^+^]_*E*_ increases, the amplitude of oscillation diminishes. This is also consistent with the experimental results in [48].

**Fig 7.**
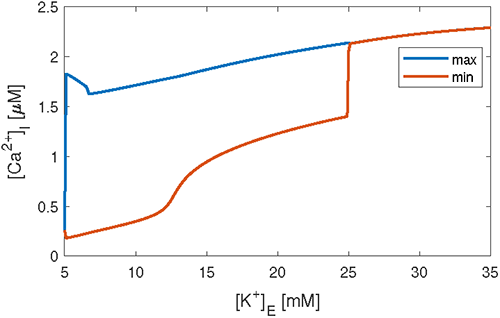
Bifurcation diagram for [*Ca*^2+^]_*I*_ when [*K*^+^]_*E*_ vary from 5 *mM* to 35 *mM*. The critical value of [*K*^+^]_*E*_ for inducing depolarization is 27.5 *mM*.

#### Spatially varied ECS potassium

Next we consider a case with diffusion in a 2-D square region Ω_*q*_ = [0, 20] × [0, 20]. Initially, the distribution of KCl in ECS is a Gaussian with peak value 80 *mM* at the center of Ω_*q*_ (see Figure 8 (a)). Non-flux boundary conditions are imposed on each boundary of Ω_*q*_. The snapshots of [*Ca*^2+^]_*I*_ at different time steps are shown in Figure 9. It can be seen that a wave is initiated at the center of the domain where [*K*^+^]_*E*_ is the highest and propagate around into low [*K*^+^]_*E*_ region. Figure 8 (b) shows [*Ca*^2+^]_*I*_ dynamics of several points on the line *x*_2_ = 10 in Ω_*q*_. It shows that the oscillation is propagated from the stimulus center to the non-stimulus far region with varied frequencies. While, the center point keeps depolarization due to high [*K*^+^]_*I*_. The propagation is also caused by propagation of membrane potential, since the diffusion effect of [*K*^+^]_*E*_ is too small and the concentration of potassium in the non-stimulus region is lower for both “connected-connected” and “connected-disconnected” cases as shown in Figure 21 in Appendix D. Similar to Figure 4, the dynamics of ICS calcium [*Ca*^2+^]_*I*_ and ⟦*ϕ*⟧ for a single “row” (*x*_2_ = 10) of Ω_*q*_ are presented in Figure 22 in Appendix D. For the “connected-disconnected” case, the oscillation is only observed near the center of Ω_*q*_. While for the “connected-connected” case, the oscillation wave could be observed in the whole Ω_*q*_ from high [*K*^+^]_*E*_ region to low [*K*^+^]_*E*_ region for the cases both with and without ions diffusion effect.

**Fig 8.**
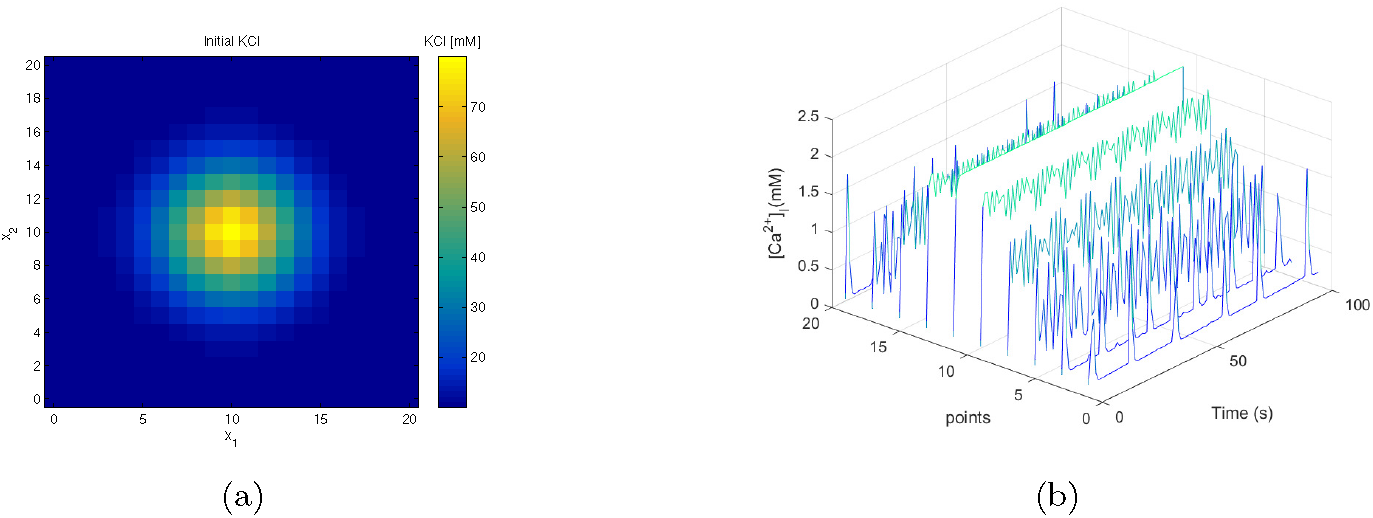
(a) Initial ECS KCl distribution. (b) [*Ca*^2+^]_*I*_ dynamics on the line *x*_2_ = 10.

**Fig 9.**
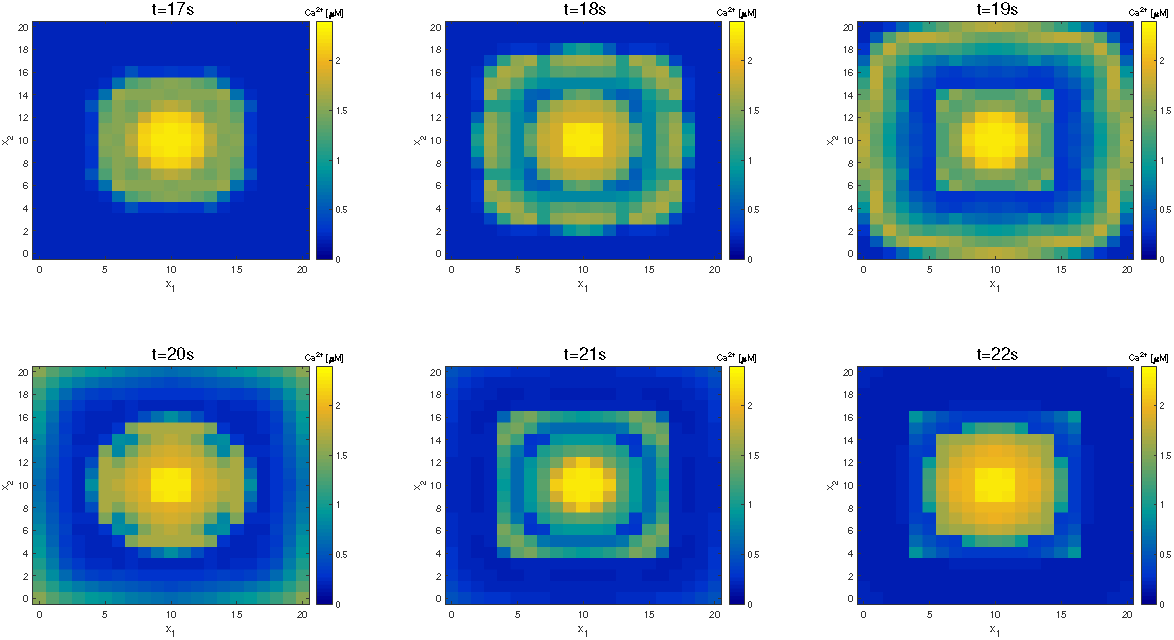
Snapshots of [*Ca*^2+^]_*I*_ at various timesteps of the 2-D square domain. A wave is initiated at the center of the domain where [*K*^+^]_*E*_ is the highest and spreads around.

**Fig 10.**
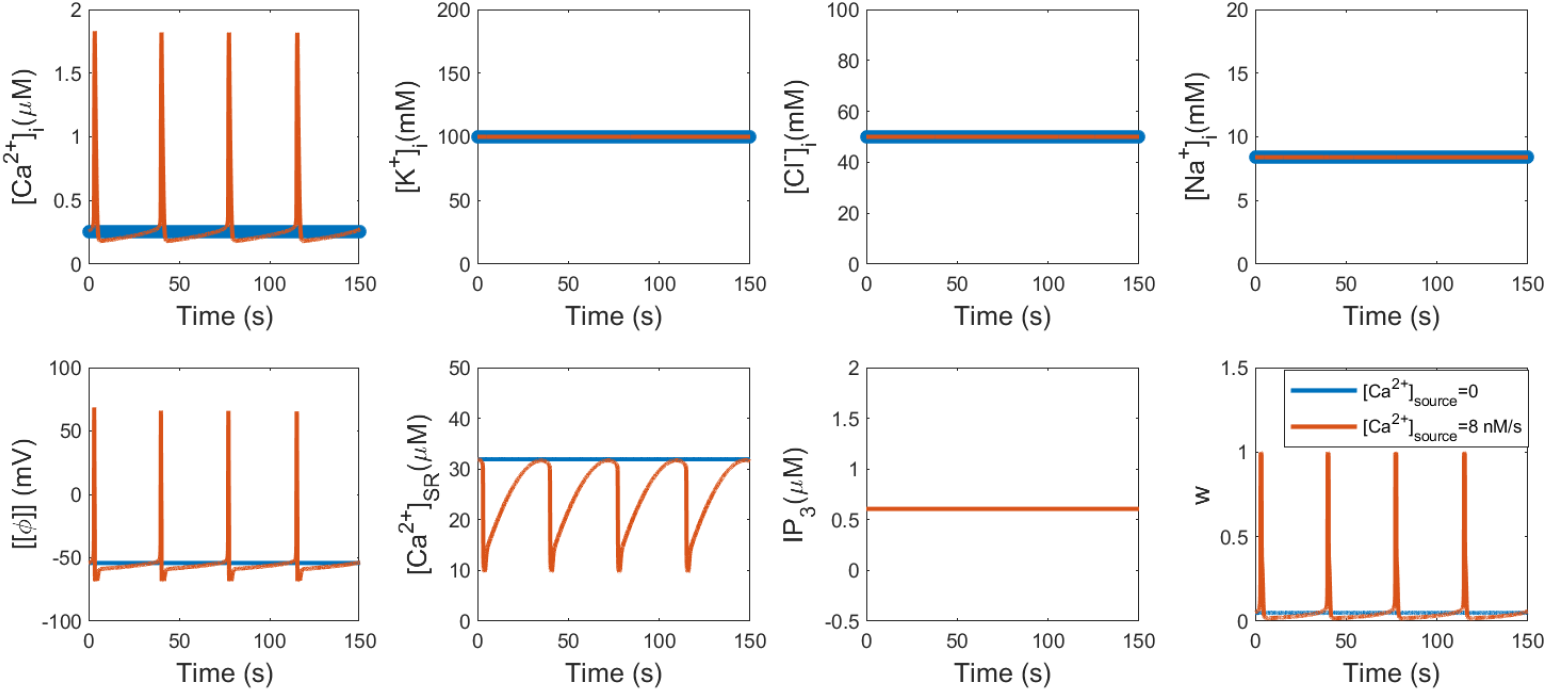
Dynamics of [*Ca*^2+^]_*I*_ with different internal *Ca*^2+^ sources for the case of space homogeneous problem. Blue lines: none source; Red lines: 8*nM/s*.

**Fig 11.**
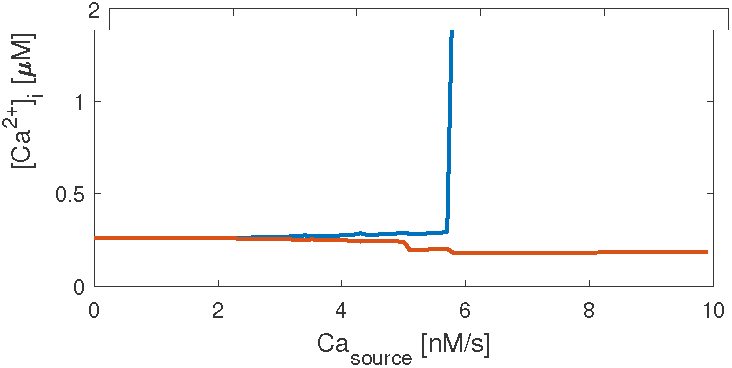
Bifurcation diagram for [*Ca*^2+^]_*I*_ when the internal *Ca*^2+^ source varies from 0 *nMs*^−1^ to 12 *nMs*^−1^.

**Fig 12.**
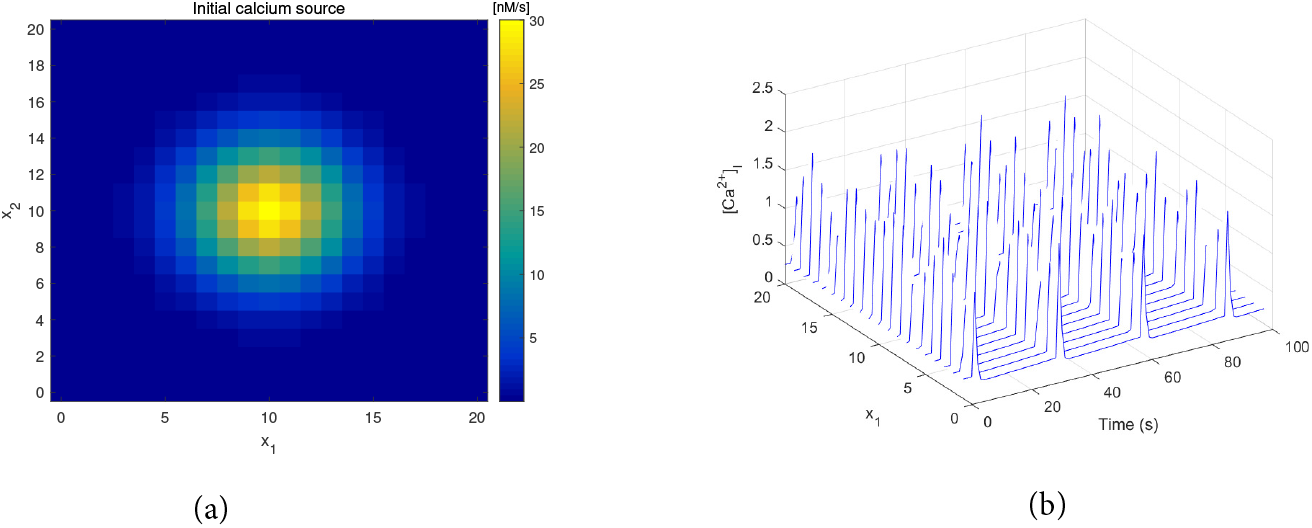
(a): Initial cons(taa)nt [*Ca*^2+^]_*I*_ source distribution; (b): [*Ca*^2+^]_*I*_ d(by)namics of points on the line *x*_2_ = 10.

**Fig 13.**
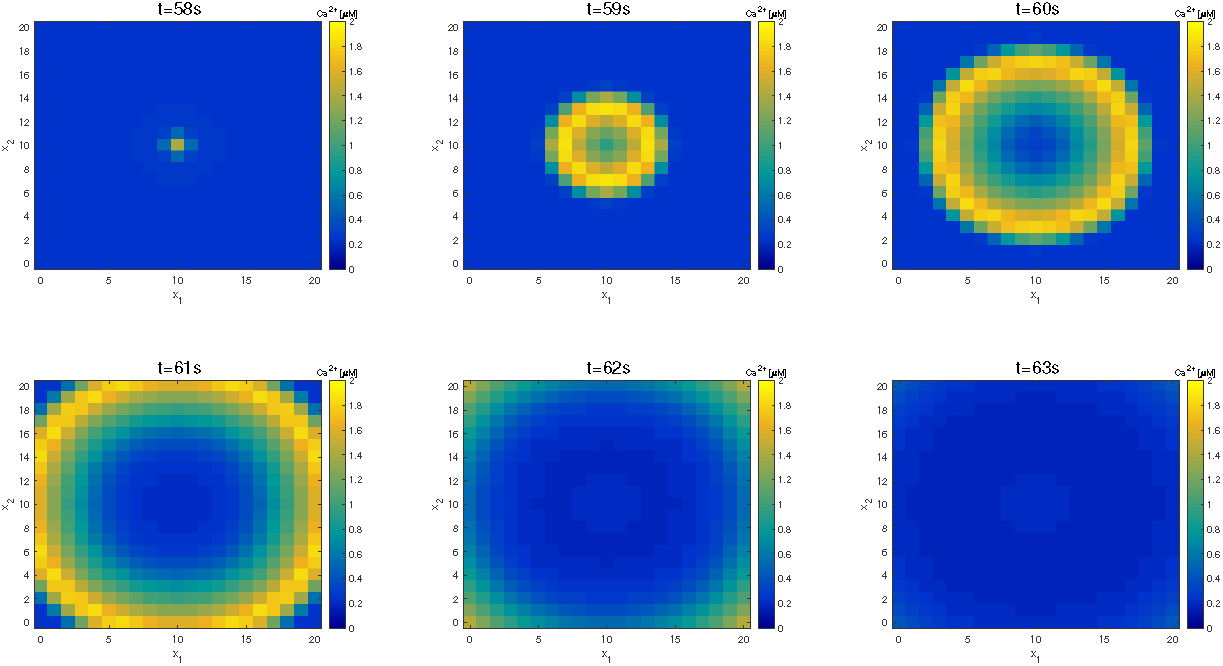
Snapshots at various timesteps of the 2-D square domain. A wave is initiated at the center of the domain where constant [*Ca*^2+^]_*I*_ source is the highest and spreads around.

**Fig 14.**
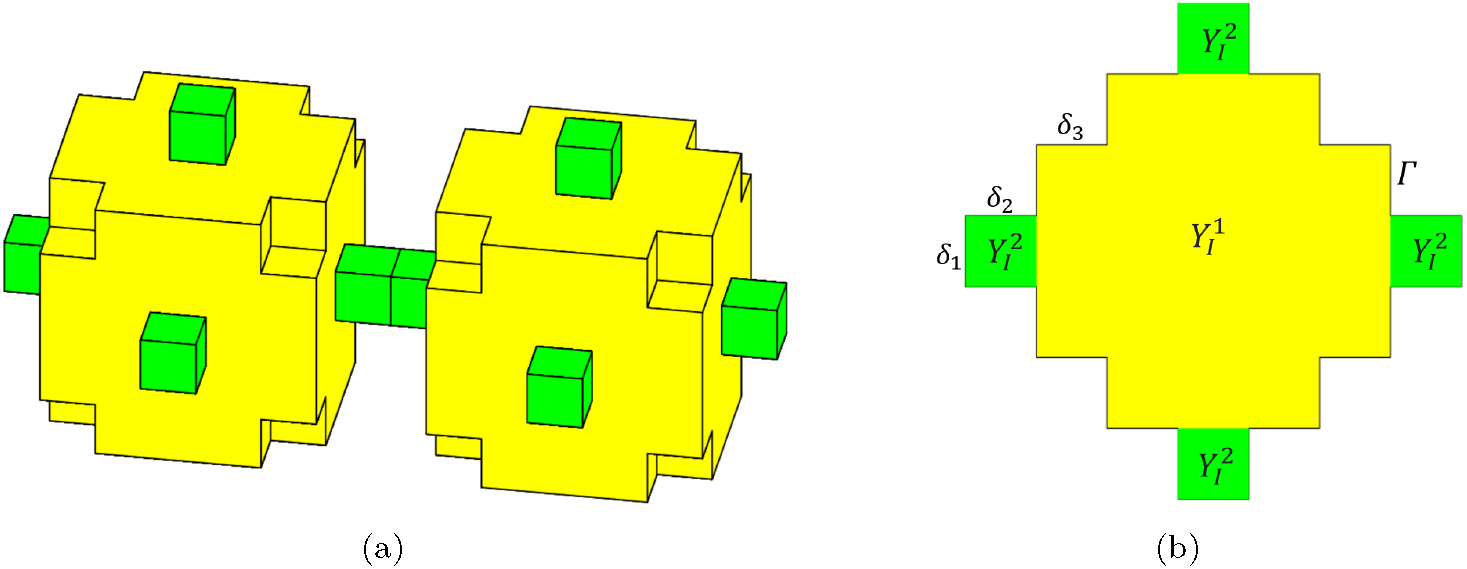
(a): 3-dimensional structure of reference cell *Y*, the green part is the gap junction, the yellow part is the cytosol; (b): The projection of one reference cell, 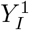 and 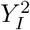 are cytosol and gap junction respectively, *δ*_1_, *δ*_2_ and *δ*_3_ are geometry parameters.

**Fig 15.**
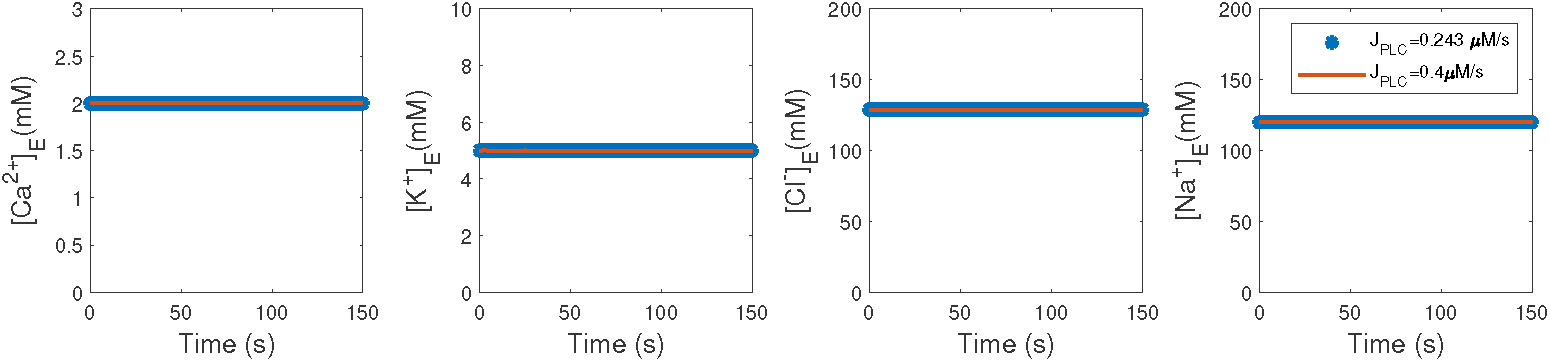
With different *J*_*PLC*_, extracellular variable dynamics for the case of space homogeneous problem.

**Fig 16.**
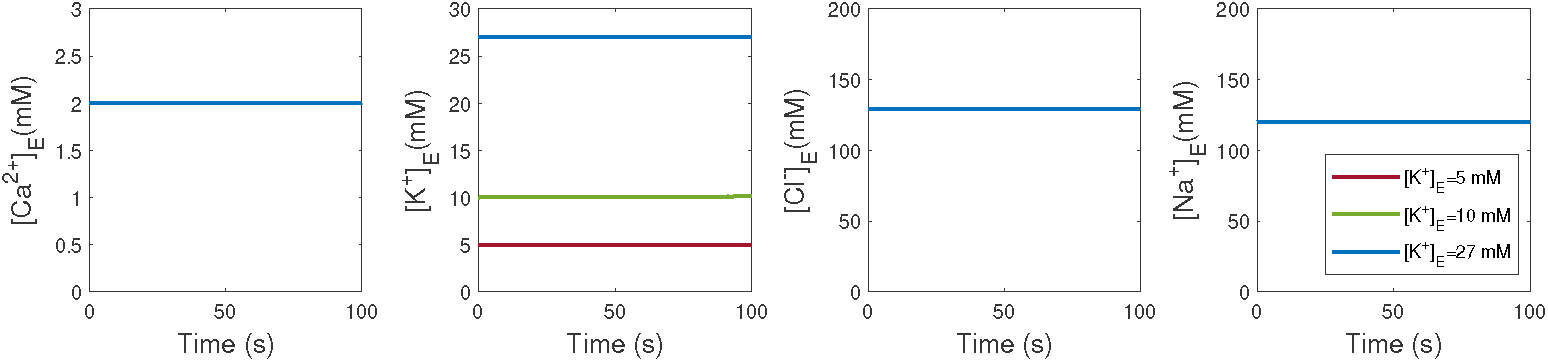
With different [*K*^+^]_*E*_, ECS variable dynamics for the case of space homogeneous problem.

**Fig 17.**
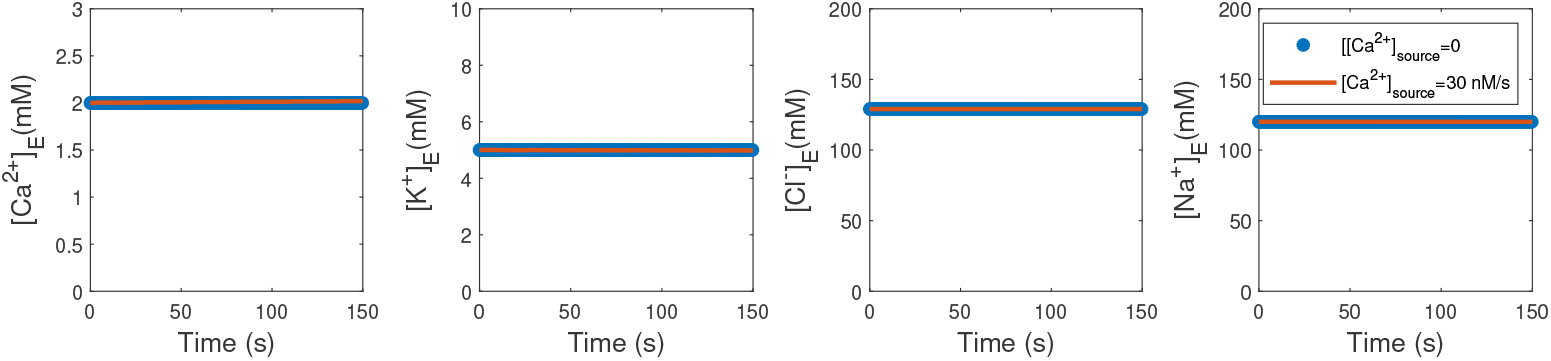
With different internal *Ca*^2+^ source, extracellular variable dynamics for the case of space

**Fig 18.**
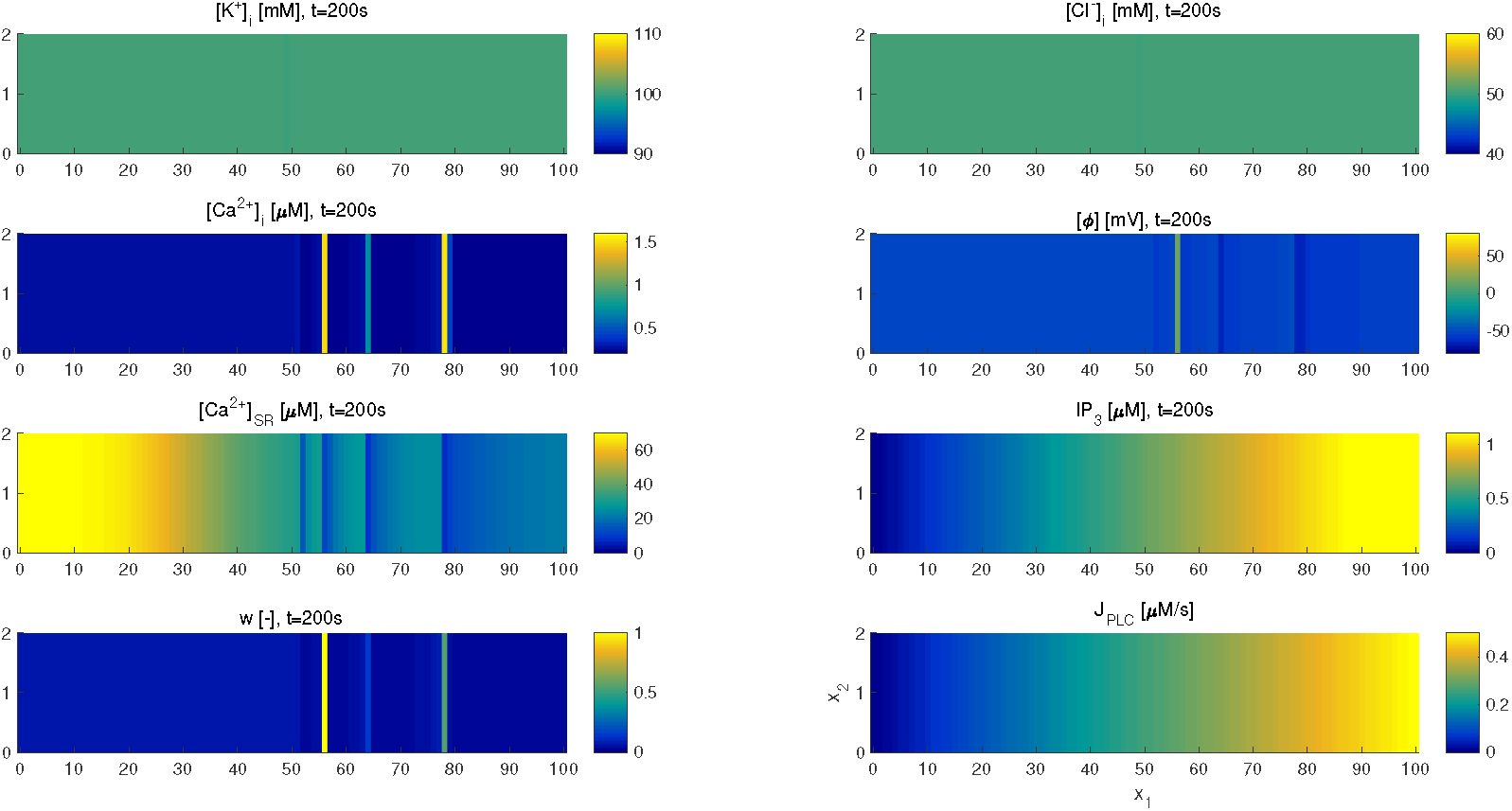
Snapshots at 200 sec of intracellular variables in Ω_*r*_ where *x*_1_ ∈ [0, 100] and *x*_2_ ∈ [0, 2], with spatially varied *J*_*PLC*_ and no diffusion.

Figure 18 and Figure 19 in Appendix D show the snapshots of ICS variables’ spatial distribution at *t* = 200 sec for the cases “connected-disconnected” and “connected-connected”, respectively. However, for the region *x*_1_ *<* 48, the results show that the *J*_*PLC*_ (or *IP*_3_ concentrations) in both case are lower than the threshold. By checking the diffusion term in (12)-(14), the scale of dimensionless effective macroscale diffusion coefficients are 10^−7^ which plays a minor role. Then what is the main mechanism to induce the oscillation in the “connected-connected” case? It is the propagation of membrane potential since the connected smooth muscle cells form a syncytium. Even though the diffusion term is tiny for ions transport, but in electric potential equations (12)_5_ and (13)_5_, the dimensionless capacitance coefficient *P*_*m*_ is of the scale 10^−5^, which amplified the effect of the diffusion of electric potential. This could be confirmed by results in Appendix D Figure 20, which compared the right hand side fluxes for electric potential equation. Another numerical test show that the phenomenon of propagation of oscillation still happens even if we eliminate the diffusion effect terms in ions transportation, while the diffusion terms in (12)_5_, (13)_5_ are preserved. The result is shown in Figure 4(c,f). It can be seen that oscillation still propagate from high *J*_*PLC*_ area to low *J*_*PLC*_ area. This verified the important role of the syncytium on the propagation of electric potential.

**Fig 19.**
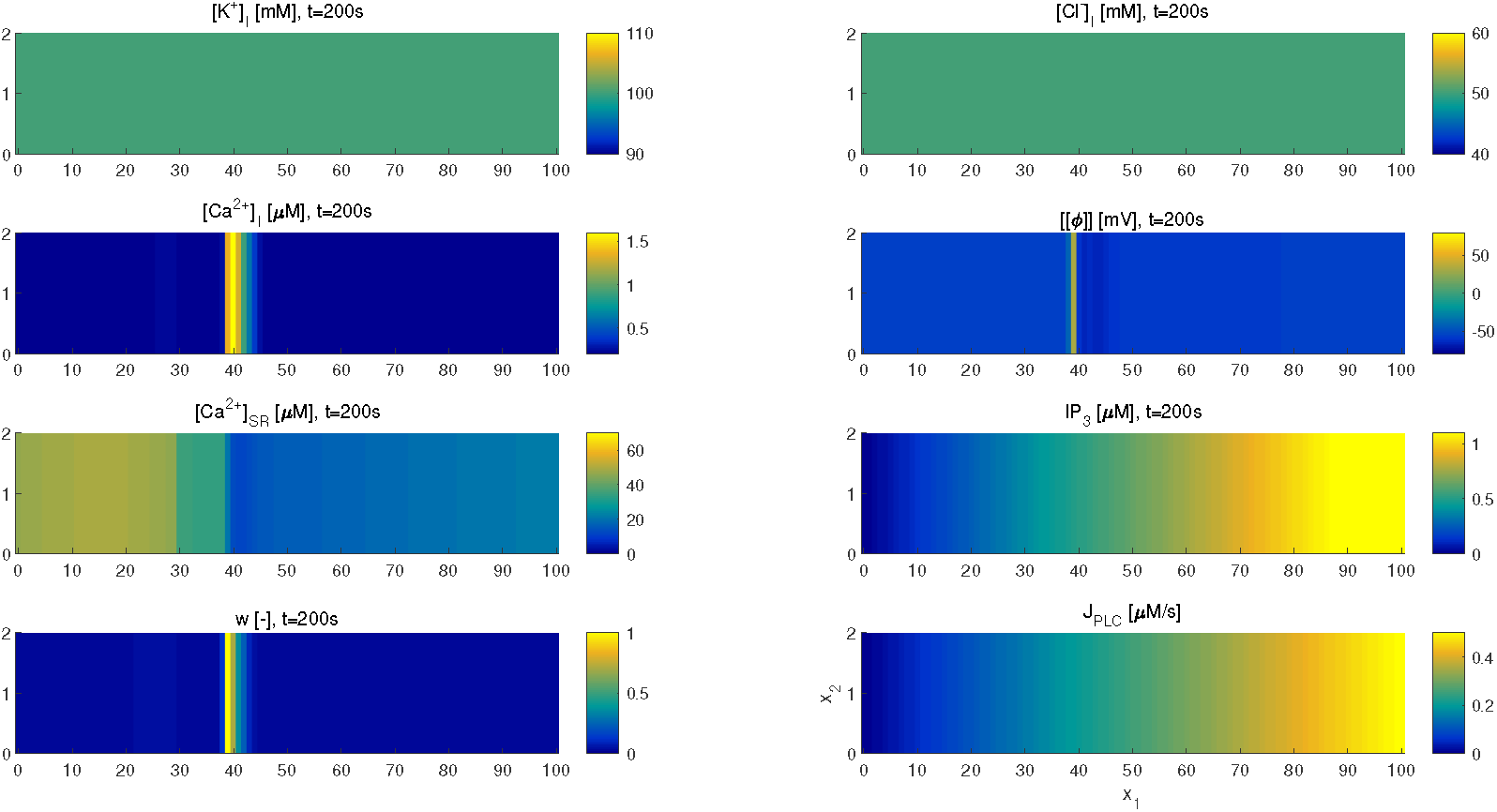
Snapshots at 200 sec of intracellular variables in Ω_*r*_ where *x*_1_ ∈ [0, 100] and *x*_2_ ∈ [0, 2], with spatially varied *J*_*PLC*_ and with diffusion.

**Fig 20.**
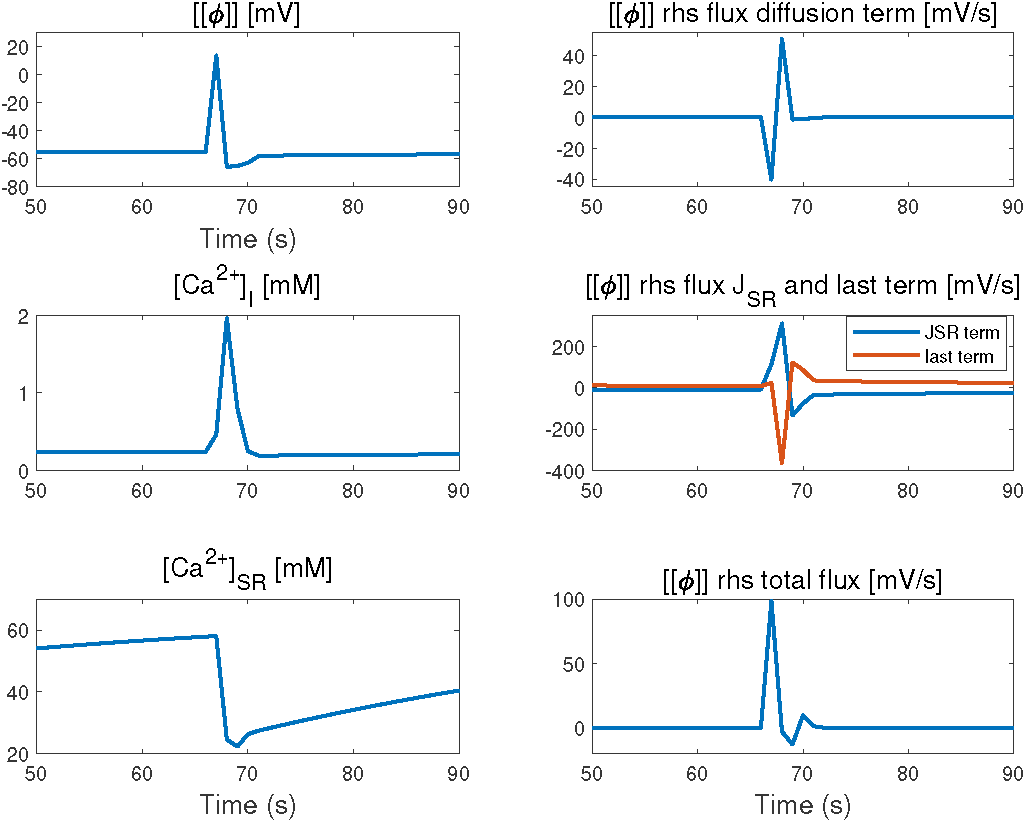
Dynamics of variables and flux with local higher *J*_*PLC*_ stimulus in “connected-connected” case. Left column: the membrane potential, concentrations of ICE and SR calcium. Right column: right hand side fluxes of electric potential equation (12)_5_ in one oscillation at the point (20,1) in Ω_*r*_.

**Fig 21.**
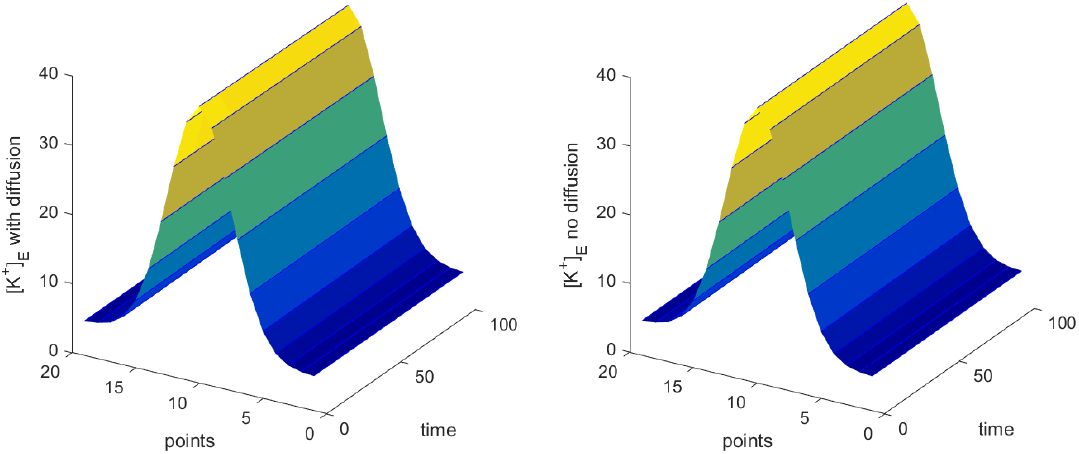
[*K*^+^]_*E*_ dynamics on the line *x*_2_ = 10 in Ω_*q*_. Left: “connected-connected” case; Right: “Connected-disconnected” case.

**Fig 22.**
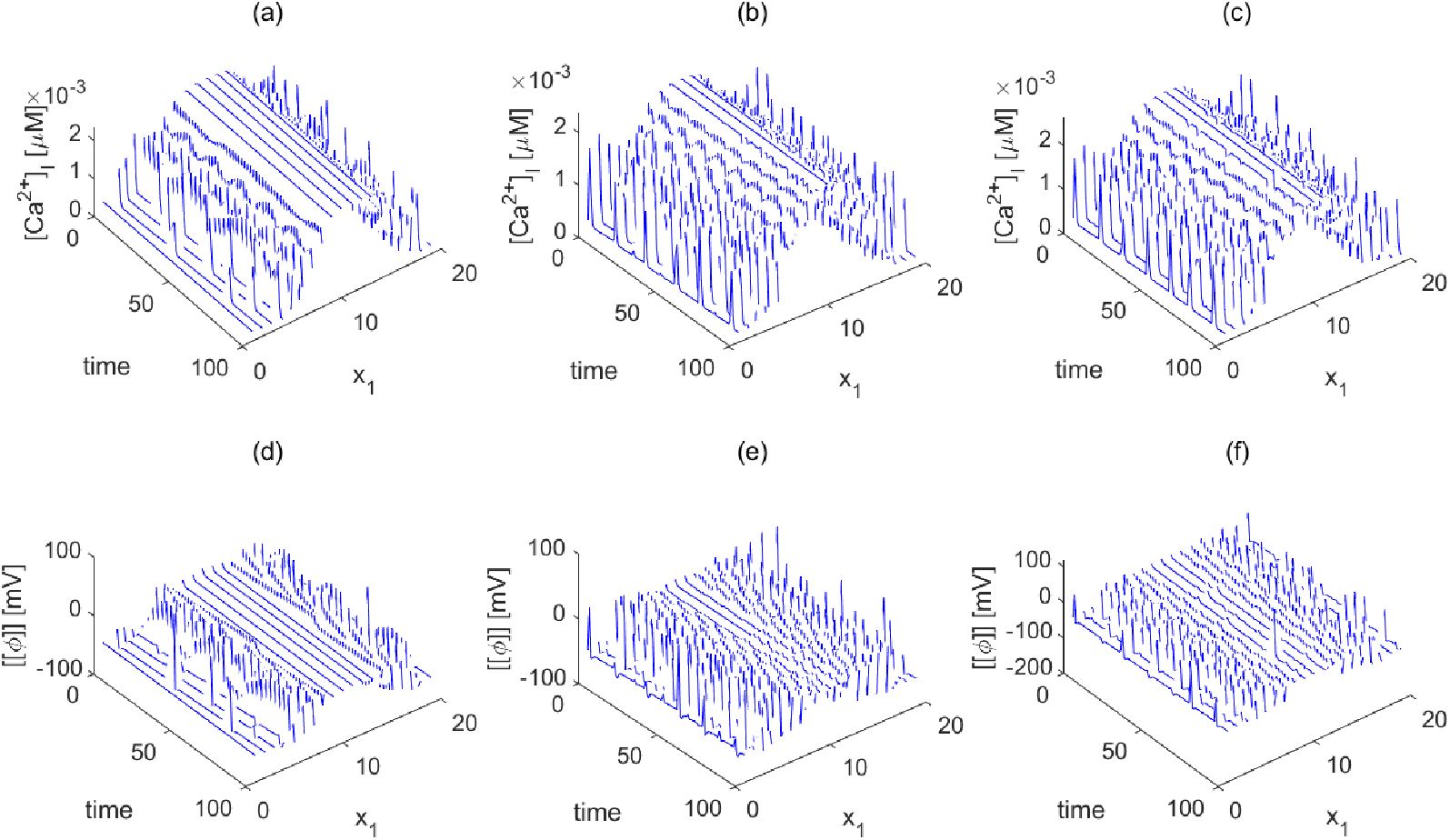
Dynamics of [*Ca*^2+^]_*I*_ and ⟦*ϕ*⟧ for a single “row” (*x*_2_ = 10) of Ω_*q*_. (a) and (d): connected-disconnected case; (b) and (e): connected-connected case. (c) and (f): connected-connected case without ion diffusion.

### Model response to varying internal *Ca*^2+^ source

In this subsection we consider the case when there is a constant internal *Ca*^2+^ source in SMCs. Migraine is related to spreading depression (SD), which is a neurophysiological phenomenon characterized by abrupt changes in intracellular ion gradients [49]. SD could cause a transient increase in light transmittance in the dendritic regions of both CA1 and the dentate gyrus, and a [*Ca*^2+^]_*I*_ increase is observed to precede this increase in light transmittance [50]. This implies that [*Ca*^2+^]_*I*_ increase is related to SD. So we add an internal constant *Ca*^2+^ source in our model and see how it will affect the system.

#### Single cell dynamics

We first add spatial uniform internal *Ca*^2+^ source. The result is shown in Figure 10. It is clearly seen that as the constant internal *Ca*^2+^ source increases from 0 (blue line) to 8*nM/s* (red lines), oscillations in ICS [*Ca*^2+^]_*I*_, SR [*Ca*^2+^]_*SR*_, membrane potential ⟦*ϕ*⟧ are observed. While for the concentration of ICS potassium, chloride, sodium and IP3, there is no observable oscilation. Figure 17 showed the dynamics of ECS variables when *Ca*^2+^ source is 0 *nMs*^−1^ and 30 *nMs*^−1^, we can see that they do not change in both cases.

In order to see the critical value of the source for intracellular *Ca*^2+^ to oscillate, a bifurcation diagram is shown in Figure 11 and the critical value of *Ca*^2+^ source is 2.1 *nMs*^−1^.

### 0.0.1 Spatially varied [*Ca*^2+^]_*I*_ source

A similar case with diffusion as in subsection is considered in Ω_*q*_ = [0, 20] × [0, 20] with spatially varied [*Ca*^2+^]_*I*_ source. Initially, the distribution of constant [*Ca*^2+^]_*I*_ source is a Gaussian with peak value 30 *nM/s* at the center of Ω_*q*_ (see Figure 12). Non-flux boundary conditions are imposed on each boundary of Ω_*q*_. In Figure 13, some snapshots of time steps. It can be seen that a wave is initiated at the center of the domain where constant [*Ca*^2+^]_*I*_ source is the highest and propagate around into low constant [*Ca*^2+^]_*I*_ source region. The dynamics of ICS calcium [*Ca*^2+^]_*I*_ and ⟦*ϕ*⟧ for a single “row” (*x*_2_ = 10) of Ω_*q*_ are presented in Figure 23 in Appendix D. For the “connected-disconnected” case, the oscillation is only observed around the center of Ω_*q*_. While for the “connected-connected” case, the oscillation is observed in the whole domain. The wave propergates from high [*Ca*^2+^]_*I*_ source region to low [*Ca*^2+^]_*I*_ source region with or without ions diffusion effect.

**Fig 23.**
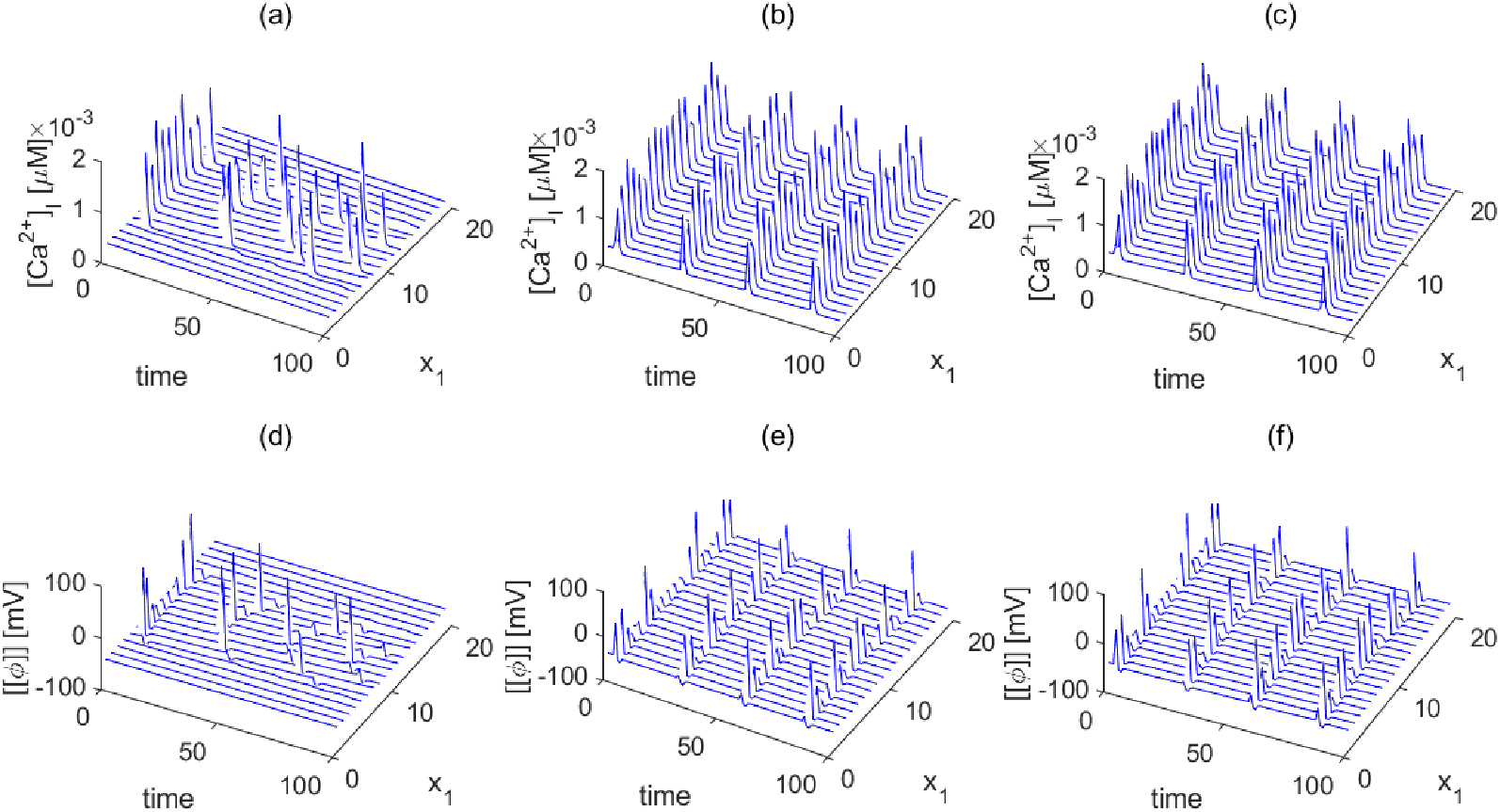
Dynamics of [*Ca*^2+^]_*I*_ and ⟦*ϕ*⟧ for a single “row” (*x*_2_ = 10) of Ω_*q*_. (a) and (d): connected-disconnected case; (b) and (e): connected-connected case. (c) and (f): connected-connected case without ion diffusion.

## Conclusion

Ion transport plays a crucial role in maintaining cell volume, membrane electrical properties, and water circulation. Dysfunctions in ion transport can lead to various diseases, including Diabetes and Spreading Depression. The development of mathematical models for ion transport in tissues is a complex task due to the presence of multiple scales in space, multi-physics interactions, and multi-component interactions.

In the current study, we derived effective macroscale electro-neutral bi-domain ion transport models for both connected-connected and connected-disconnected cases through asymptotic expansion. These models were used to investigate the effects of different stimuli, such as increased Phospholipase C, extracellular potassium, and intracellular calcium release, on the dynamics of cytosolic calcium during spreading depression [36, 51].

We first performed space-uniform tests, simulating a single cell, to identify the bifurcation points of different stimuli. The results were consistent with existing experimental findings, confirming the validity of our model and parameters. The results also showed that extracellular variables do not exhibit significant variation when there is higher Phospholipase C or intracellular calcium source, supporting the use of models with constant extracellular variables.

Next, we used the calibrated models to study spatially varied stimuli. Both dimensionless analysis and numerical simulations showed that the propagation of oscillations from the stimulus region to the non-stimulus region was primarily driven by the electric potential propagation along the syncytium membrane, rather than the limited diffusion of ions between cells through connexins.

However, there are some limitations to our model that we acknowledge. Firstly, we only considered simple ion channels on the membranes [41], and the communication between the cytosolic and mitochondrial compartments [52] was simplified using a source term. Additionally, we only use trivial 2D domain for computation without considering the real 3D structure of smooth muscle cells in the human vasculature [53]. In the future, our model could be expanded to consider specific channels [54] and neuron-glial compartments [18], which would allow for a deeper understanding of the effects of spatial variation of stimuli on neurovascular coupling and cerebral autoregulation [55–59].

In conclusion, our study provides valuable insights into the mechanisms of ion transport during spreading depression and highlights the potential of mathematical models in understanding complex biological systems.

## Supporting information

### Asymptotic Expansion

The two different lengthscales imply that the derivatives must be transformed according to

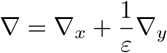

Based on the positivity assumption, it is obvious that when *ε* is sufficiently small, we have

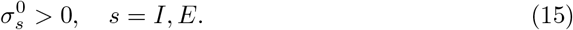

And EN condition (1) imply that

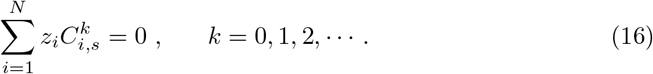

Since *J*_*i*_ is smooth, according to Taylor expansion, we have

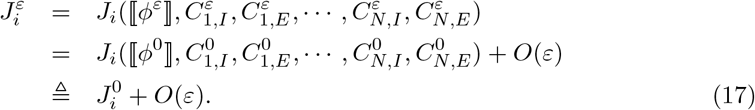

For 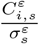 we have

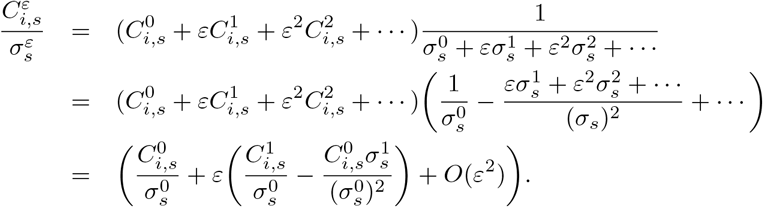

So

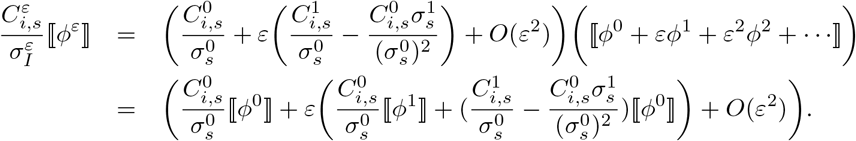

For “connected-disconnected” case and for intracellular region 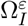, substituting the two-scale asymptotic expansions of relevant functions into (5) leads to

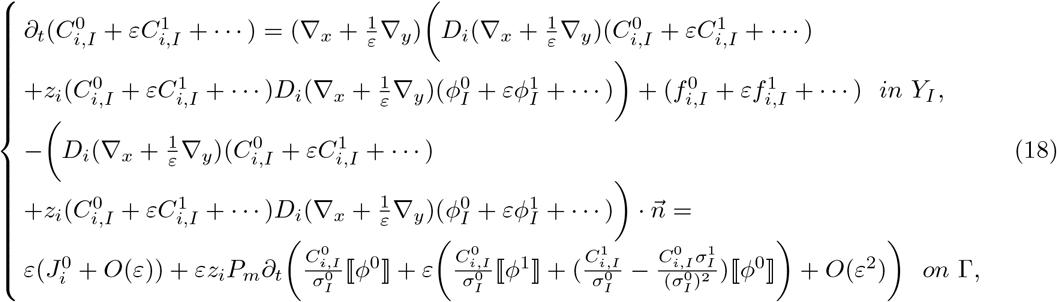

The coefficients of *ε*^−2^:

Firstly, the coefficients of *ε*^−2^ in (18)_1_ and the coefficients of *ε*^−1^ in (18)_2_ are

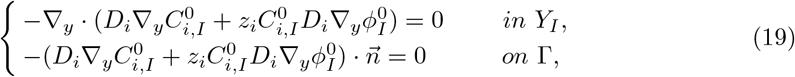

Multiply (19) by 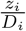 and sum for *i* = 1, …, *N*, noticing (16), it follows that

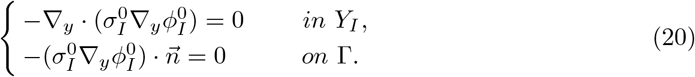

According to (15) and (20) we can deduce that 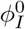 is independent of *y*, i.e.

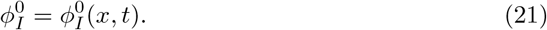

In fact, multiplying (20) by 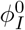 and integrating in *Y*_*I*_ yields

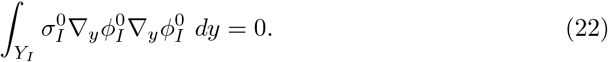

By (15), we can derive from the above equation that 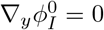, so 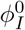 is independent of *y*.

Then by substituting (21) into (19), we find

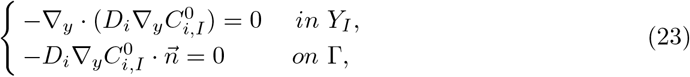

which means 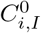 is independent of *y*, so

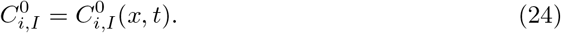

The coefficients of *ε*^−1^:

The coefficients of *ε*^−1^ in (18)_1_ and the coefficients of *ε*^0^ in (18)_2_ are

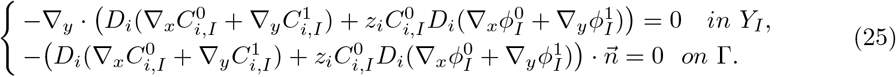

Multiply (25) by 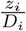 and sum for *i* = 1, …, *N*, from (16), it follows that

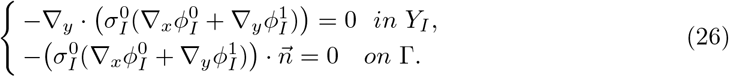

Since 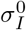 is independent of *y*, we can deduce from (26) that

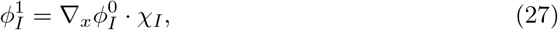

where 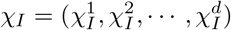, and satisfy

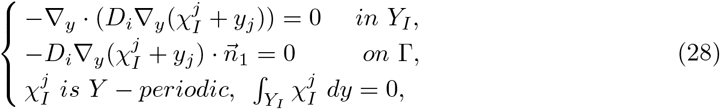

for *j* = 1,, *d*. Since *Y*_*I*_ is completely included in *Y*, we deduce that the solution of (28) is

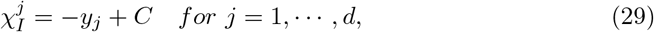

where *C* is a constant. This implies

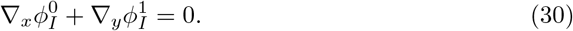

Then, by substituting (30) into (25), we obtain

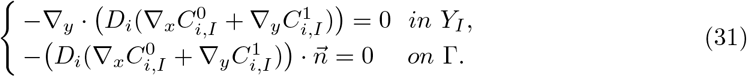

Similarly, we have

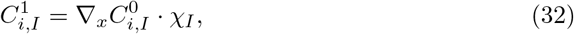

and

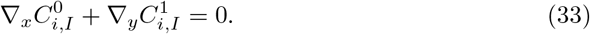

The coefficients of *ε*^0^:

The coefficients of *ε*^0^ in (18)_1_ and the coefficients of *ε*^1^ in (18)_2_ are

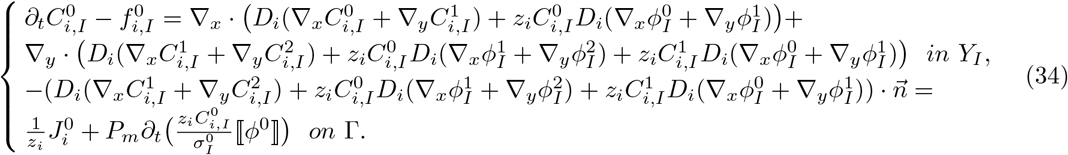

Integrating (34) in *Y*_*I*_ and noticing (30), (33), we obtain

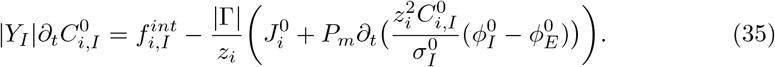

Multiply the above equation with *z*_*i*_ and summing for *i* = 1, …, *N* leads to

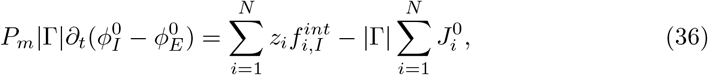

where 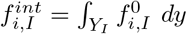.

The above two equations are homogenized equations for intracellular region 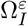, i.e., (7)_2_ and (8)_2_. The process of deriving the homogenized equations for 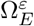 is almost the same as for 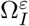. However, there are some small differences, and the resulted homogenized equations are different.

The coefficients of *ε*^−2^:

The coefficients of *ε*^−2^ for equations in 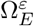 are

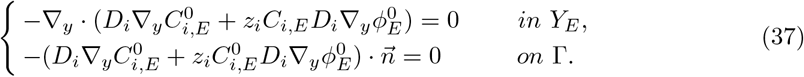

Multiply (37) by 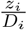 and sum for *i* = 1, …, *N*, noticing (16), it follows that

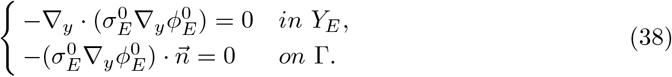

we conclude that both 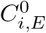 and 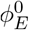 are independent of *y*, i.e.

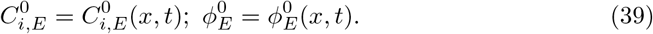

The coefficients of *ε*^−1^:

The coefficients of *ε*^−1^ for equations in 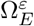 are

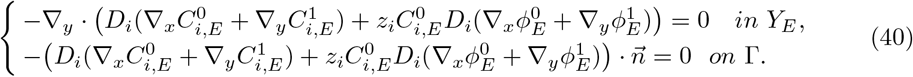

Multiply (40) by 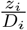 and sum for *i* = 1, …, *N*, from (16), it follows that

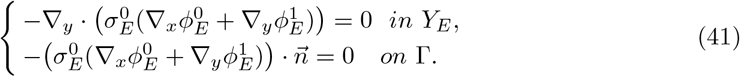

So we have

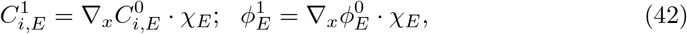

where 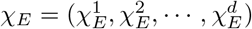, satisfy

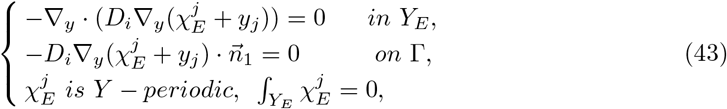

for *j* = 1, …, *d*. Here we could not derive that the solution of (43) is 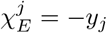, because *Y*_*E*_ reaches the boundary of *Y* in “connected-connected” case.

The coefficients of *ε*^0^:

The coefficients of *ε*^0^ for equations in 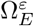 is

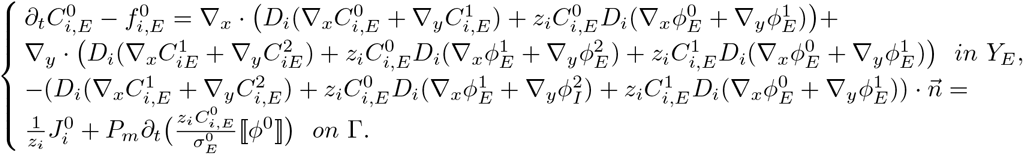

So, integrating (44) in *Y*_*E*_ leads to

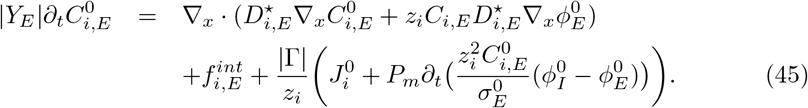

Multiply the above equation with *z*_*i*_ and summing for *i* = 1, …, *N* yields

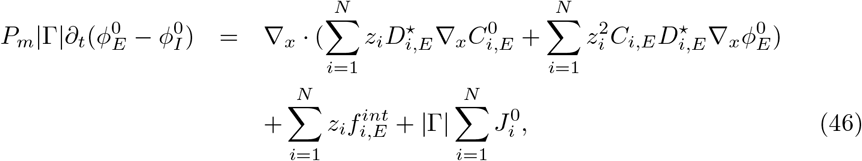

where 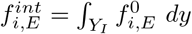 and 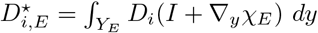. (45), (46) are (7)_1_, (8)_1_. The “connected-disconnected” case is proved.

For “connected-connected” case, the micro structure of the domain is different. In reference cell *Y*, both *Y*_*I*_ and *Y*_*E*_ reach the boundary of *Y*. The derivation of homogenized equations is quite similar to the process of “connected-disconnected” case, so we omit the proof for this case. The difference is when we consider the coefficients of *ε*^−1^ in 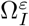, the solution 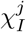 of (28) is no longer −*y*_*j*_. This is because *Y*_*I*_ reaches the boundary of *Y*. The theorem is proved.

### Effective Diffusion Estimation

Next we estimate the effective diffusion coefficient *D*^⋆^ and give the initial values for the numerical simulations in section.

Suppose the 3-dimensional geometry of one cell as Figure 14, and we can solve problem (11) to get 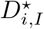. Gap junction is a specialized intercellular connection between a multitude of animal cell-types, including SMCs. They directly connect the cytoplasm of two cells, which allows various molecules, ions and electrical impulses to directly pass through a regulated gate between cells. So in the reference cell *Y*, the intracellular part *Y*_*I*_ can be divided into 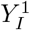 and 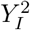, which represent the cytosol and gap junctions respectively (see Figure 14). Suppose the cell is a cube and the cross section of the gap junction is square. Another two factors needed are the geometry parameters and diffusion coefficient in 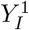 and 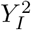. Denote by *δ*_2_ the value of the half length of the gap junction, *δ*_1_ the side length of the cross section of the gap junction (*δ*_1_, *δ*_2_ and *δ*_3_ are dimensionless values). The structure of the reference cell is uniquely determined by *δ*_1_, *δ*_2_ and *δ*_3_. To match the volume of a single medial SMC(about 1630 ± 640 *μm*^3^ [60]), the dimensional side length *l* of the cell is set to be 13*μm*. Since the gap junctions occupy about 0.29% of the SMC surface [61], *δ*_1_ is set to be 0.054. A gap junction channel is formed by end-to-end docking of two hemichannels called connexin [62], which span three regions: the ICS part, the membrane part and the space between two cells. The length of these three parts are about 1.9 *nm* [62], 7.5 ∼ 10 *nm* [63] and 2 ∼ 4 *nm* [64] respectively. Choosing the middle value, the length of the connexin is about 12.15 *nm*, so *δ*_2_ is set to be 12.15 *×* 10^−3^*/*13 = 0.9346 *×* 10^−3^. To match the ECS volume fraction |*Y*_*E*_|(which is set to be 0.25), *δ*_3_ is set to be 0.3126. Since we consider the 3-dimensional case and the value of *δ*_2_ is too small, it is numerically expensive to compute 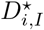. To estimate 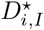, we fix the value of *δ*_1_, *δ*_3_ and change the value of *δ*_2_ as 2*/*72, 3*/*72, …, 10*/*72, and the values of 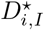 are computed respectively. Eventually we can apply curve fitting to estimate 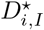.

Denote by *D*_*A*_, *D*_*B*_ the micro diffusion coefficients in 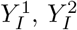 respectively. Although 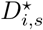 are matrix, but numerical results show that the off-diagonal elements are zero and the diagonal elements are the same, so we can simply assume that 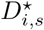 are scalars.

Using finite volume method to solve (11), we can apply 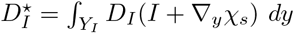 to compute the effective diffusion coefficients. The mesh size is 72 *×* 72 *×* 72.

In order to estimate 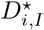, the value of *D*_*A*_, *D*_*B*_ are needed. *D*_*A*_ is the diffusion coefficient in cytosol and it can be found in references. However, since the gap junction part 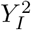 is consist of several connexin channels, *D*_*B*_ is actually an effective diffusion coefficient in the gap junction. Let *D*_*p*_ be the diffusion coefficient in the pore of each connexin channel, *D*_*aq*_ be the diffusion coefficient in water, then it is estimated that [65]

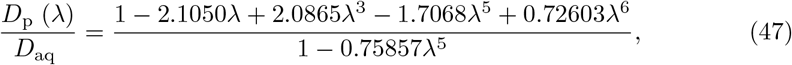

where *λ* = *R*_*ion*_*/R*_*pore*_ is the ratio of the permeant (*R*_*ion*_) to the pore (*R*_*pore*_) diameter. Usually, *R*_*pore*_ is about 1 ∼ 1.4 *nm* [66–69]. We take the middle value and let *R*_*pore*_ = 1.2 *nm*. And the value of *D*_*A*_ ranges from 30% to 90% of the corresponding value of *D*_*aq*_ [65]. So if *λ, D*_*A*_ are known, the range for *D*_*p*_ can be estimated.

The number of gap junctions in one SMC is about 145 [61], so there are about 24 on each face of *Y*. In addition, based on diffusion law, we have

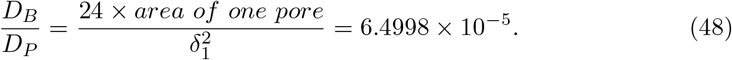

When computing the area of one pore, we used the value of *R*_*pore*_. The values of *R*_*ion*_ for different ions can be found in [70]. *R*_*ion*_ for *Ca*^2+^, *K*^+^, *Cl*^−^, *Na*^+^ are 0.204 *nm*, 0.276 *nm*, 0.362 *nm*, 0.204 *nm* respectively. We will also need *R*_*ion*_ for molecule *IP*_3_. Since the polar surface area of *IP*_3_ is about 261 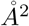, suppose the molecule is a sphere, then *R*_*ion*_ for *IP*_3_ can be estimated to be 0.911 *nm*. The dimensional micro diffusion coefficients *D*_*A*_ for *Ca*^2+^, *K*^+^, *Cl*^−^, *Na*^+^ are [22, 71]: *D*_*Ca*_ = 2.23 *×* 10^−10^ *m*^2^*/s, D*_*K*_ = 1.96 *×* 10^−9^ *m*^2^*/s, D*_*Cl*_ = 2.03 *×* 10^−9^ *m*^2^*/s, D*_*K*_ = 1.33 *×* 10^−9^ *m*^2^*/s*, 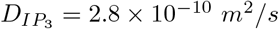 respectively. Then from the value of 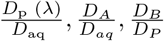 we can compute *D*_*B*_. Now, based on curve fitting, 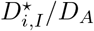 can be estimated for different ions (see Table 4).

**Table 4.** 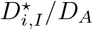 for different ions and *IP*_3_

| ion | $Ca^{2+}$ | $K^+$ | $Cl^-$ | $Na^+$ | $IP_3$ |
| --- | --- | --- | --- | --- | --- |
| $D_{i,I}^*/D_A$ | $4.757 \times 10^{-5}$<br>$\sim 1.427 \times 10^{-4}$ | $3.904 \times 10^{-5}$<br>$\sim 1.171 \times 10^{-4}$ | $3.029 \times 10^{-5}$<br>$\sim 9.086 \times 10^{-5}$ | $4.711 \times 10^{-5}$<br>$\sim 1.413 \times 10^{-4}$ | $2.088 \times 10^{-6}$<br>$\sim 6.264 \times 10^{-6}$ |

For 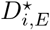, according to [72], we have

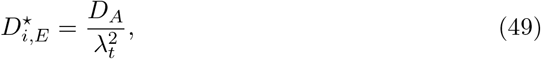

where *λ*_*t*_ ≈ 1.6 [73] is the tortuosity factor for ECS.

There are negatively charged protein molecules (*A*^−^) manufactured inside the cell, suppose the valence of these molecules is −1.5 and the concentration is [*A*^−^]_*I*_.

### Definition of Flux

Fluxes in (12), (14), (13) are (see [74], [39], [75]).

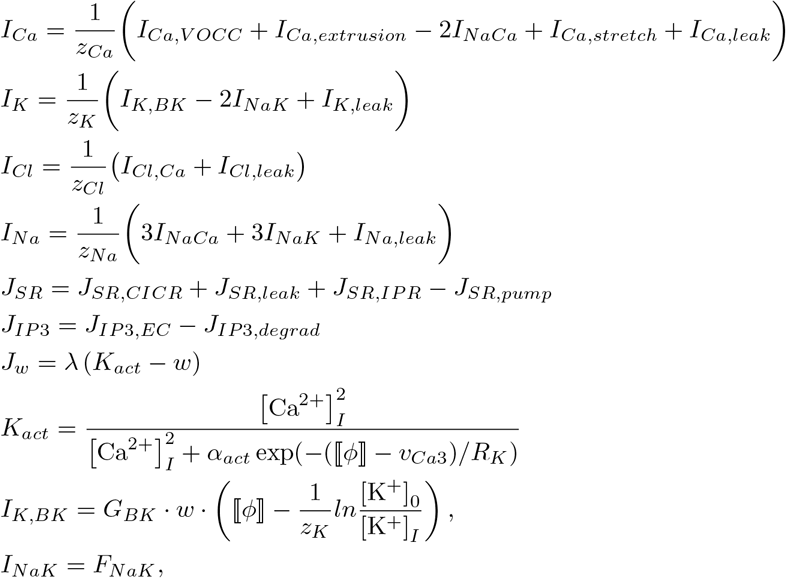

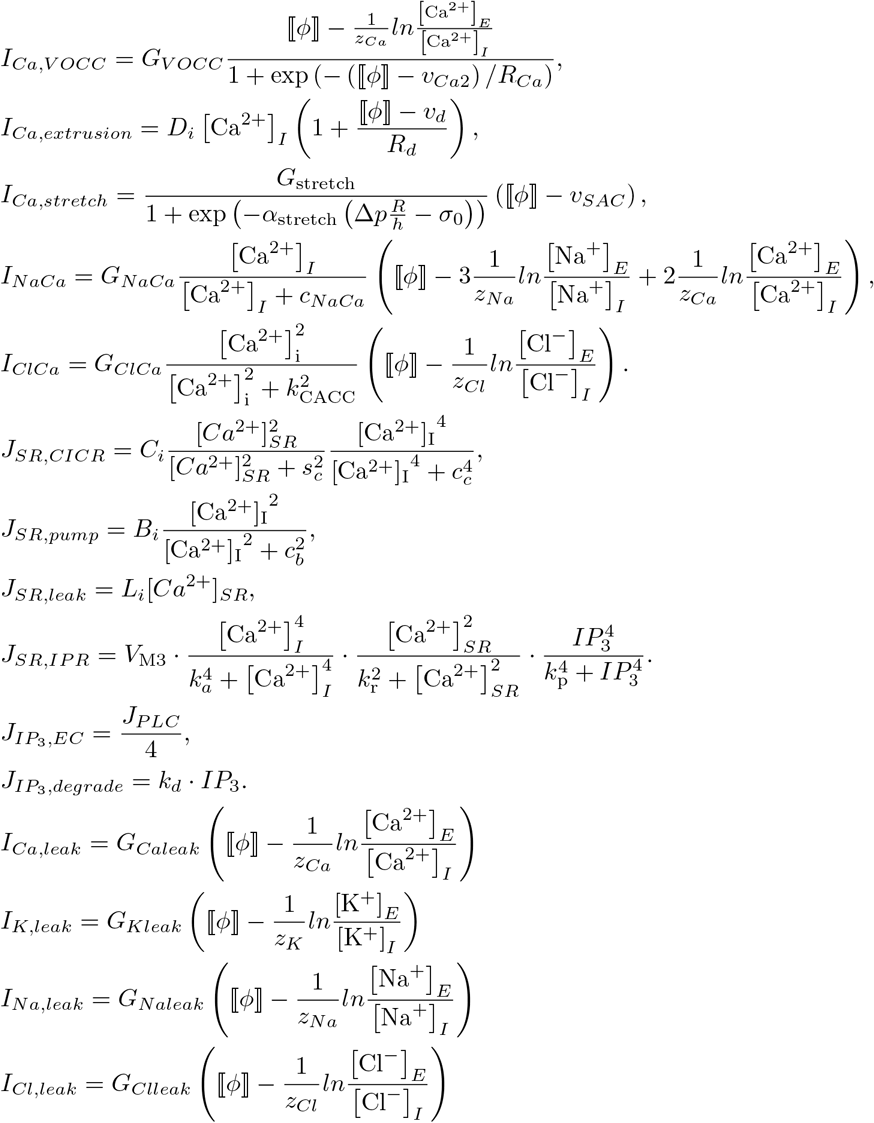

Initially, the fluxes are set to be zero to reach an equilibrium state, and the leak terms are added to help reach this equilibrium.

## Acknowledgments

This work was partially supported by the National Natural Science Foundation of China (12071190, 11971342, 12231004), Natural 11 Science Foundation of Guangdong Province of China (2023A1515010803), the Fundamental Research Funds for the Central Universities (2042021kf0050) and NSERC (CA) (RGPIN-2016-05306). The first author is grateful for the financial support from China Scholarship Council (CSC) during his visit to York University in Toronto, where this work is partially done.

